# Dynamic traction force measurements of migrating immune cells in 3D matrices

**DOI:** 10.1101/2022.11.16.516758

**Authors:** David Böhringer, Mar Cóndor, Lars Bischof, Tina Czerwinski, Niklas Gampl, Phuong Anh Ngo, Andreas Bauer, Caroline Voskens, Rocío López-Posadas, Kristian Franze, Silvia Budday, Christoph Mark, Ben Fabry, Richard Gerum

## Abstract

Immune cells such as natural killer (NK) cells migrate with high speeds of several µm/min through dense tissue, but the traction forces are unknown. We present a method to measure dynamic traction forces of fast migrating cells in non-linear biopolymer matrices. The method accounts for the mechanical non-linearity of the 3D tissue matrix and can be applied to time series of confocal or bright-field image stacks. The method is highly sensitive over a large range of forces and object sizes, from ∼1 nN for axon growth cones up to ∼10 µN for mouse intestinal organoids. We find that NK cells display bursts of large traction forces that increase with matrix stiffness and facilitate migration through tight constrictions.

## Introduction

Essential cell functions such as migration and spreading require that cells exert traction forces on their extracellular matrix. Several traction force microscopy methods exist that measure the force-induced deformations of the extracellular matrix, and reconstruct the cell forces based on continuum-mechanical principles. Mathematically, the computation of forces from matrix deformations is an inverse problem where small measurement noise can give rise to large erroneous forces. This is typically solved by various smoothing or force regularization methods (1–3). For cells grown on two-dimensional (flat) matrices with linear elastic properties, such as polyacrylamide hydrogels or silicone elastomers, robust and computationally inexpensive methods have been developed (4–7). The force reconstruction problem becomes considerably more difficult in 3 dimensions (2, 8–11), especially in the case of mechanically non-linear materials, such as collagen or fibrin hydrogels (12–18), which are frequently used as a 3D matrix for cell culture.

Current 3D force reconstruction methods typically require the recording of a large 3D image stack around a cell that is sufficiently far away from other cells. In addition, knowledge of the cell surface (Fig. 1a top), where by definition the traction forces are located, is also required for most methods. Staining and 3D imaging the cell surface, however, can affect cell behavior or cause photo-damage. All current methods require knowledge of the force-free reference configuration of the matrix, which is obtained either by detaching the cell from the matrix, by the addition of drugs that suppress cell forces (e.g. cytochalasin D), or by killing the cell, e.g., with high-intensity laser light (Fig. 1b top). Such methods therefore pose limitations for measuring traction forces of rapidly moving cells, such as immune cells, or for investigating multiple cells within a culture dish.

**Fig. 1.**
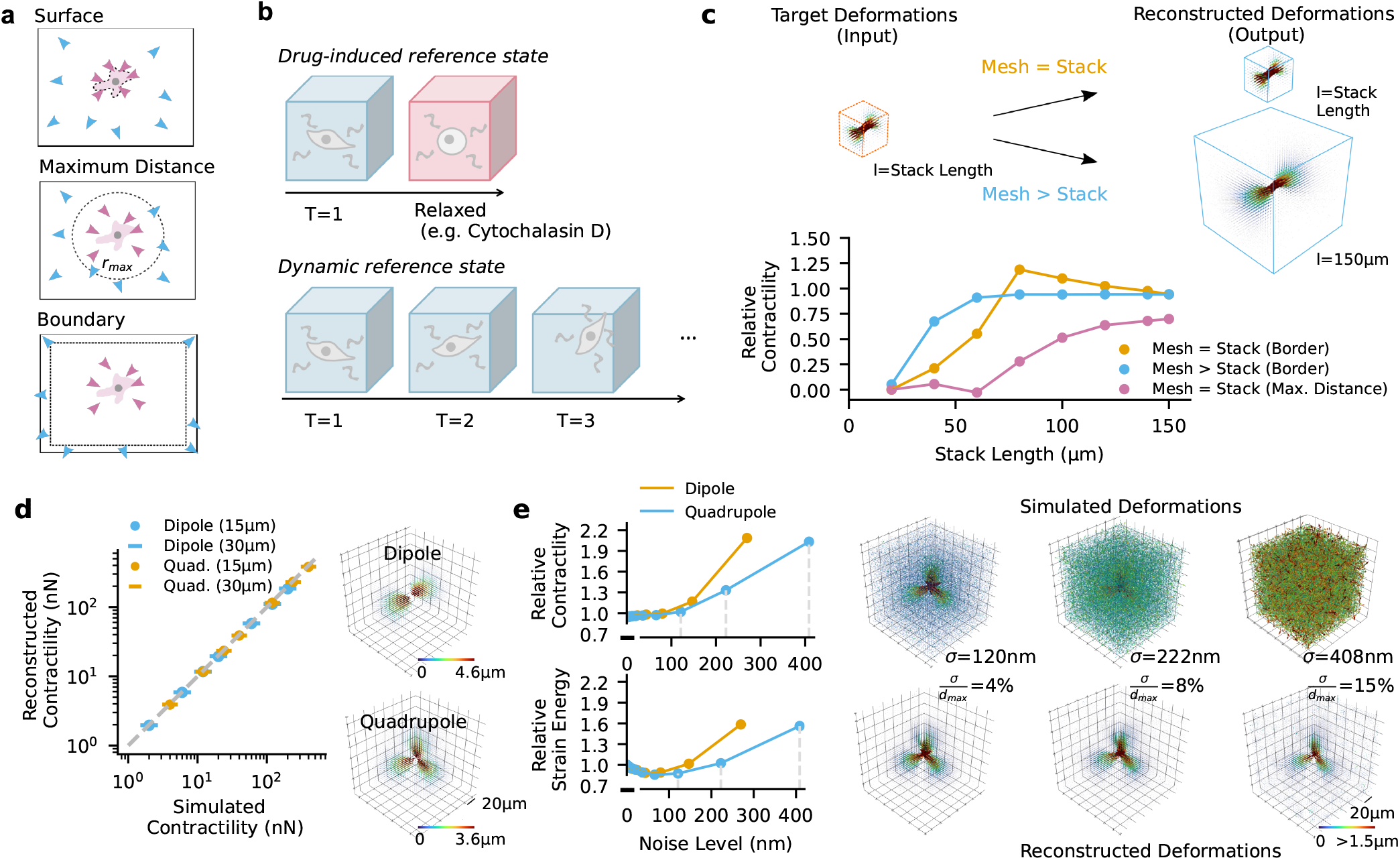
Material model: **a**, Schematic of different regularization strategies. Top: Traction force microscopy (TFM) typically constrains forces (purple) to the cell surface. Middle: Body force microscopy with global regularization reconstructs forces without knowledge of the cell surface. Forces within a defined distance (*r*_max_) around the force epicenter are considered cell-generated forces (purple), forces beyond *r*^*max*^ (dotted gray line) are considered balancing forces (blue) that also account for noise, drift, or contractile cells outside the field of view. Bottom: Saenopy is a body force microscopy method where forces are regularized throughout the image stack except for the boundary regions. This ensures that balancing forces appear only at the boundaries. **b**, Schematic of two methods for measuring the force-free matrix configuration. Top: Cell forces are relaxed using actin-depolymerizing or myosin-inhibiting drugs. 3D matrix deformations are measured from two image stacks recorded before and after drug treatment. Bottom: The force-free matrix configuration is computed from the cumulative matrix displacements between consecutive image stacks of a time series, median-averaged over the observation period. Dynamic 3D matrix deformations are obtained from the cumulative matrix displacements relative to the median average. **c**, Influence of image stack size and reconstruction volume on force reconstruction accuracy. The cell is simulated by a force dipole with 20 nN and a distance of 30 μm. The image stack size (cube length) is increased between (20 μm-150μm)^3^. The reconstruction volume (mesh) has either the same dimensions as the image stack (yellow) or has a constant size of (150 μm)^3^ (blue). Force reconstruction is performed with Saenopy regularization (yellow and blue lines) or with global regularization (*r*_max_ = 50 μm, magenta line). **d**, Accuracy of force reconstruction for simulated dipoles (upper inset) and quadrupoles (lower inset) for different contractilities. Dipoles or quadrupoles are simulated as pairs of monopoles either 15 μm or 30 μm apart. Contractility is the sum of the force monopoles. Dashed gray line shows line of identity. **e**, Influence of noise on force reconstruction (top) and strain energy (bottom) for simulated dipoles (contractility of 20 nN) or quadrupoles (contractility of 40 nN). Maximum matrix deformations are around 3 μm in both cases. Noise is added as Gaussian noise with specified standard deviation *σ* to the deformation field (upper right images for three selected noise values) and is expressed as absolute sd noise σ or percentage of sd noise relative to the maximum matrix deformation 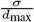. Lower right images show reconstructed deformation fields.Reconstructed contractility and strain energy are expressed relative to the input values.

## Results & Discussion

Here, we overcome these limitations by a locally adjusted regularization approach, and by obtaining the force-free reference configuration of the matrix from time-lapse images. Our method builds on a previously developed non-linear semi-affine network optimizer (Saeno) (12). This method iteratively minimizes the discrepancy between measured and reconstructed matrix deformations (Fig. 1a middle) by adjusting point forces that can appear anywhere in the volume but are regularized to favor few, large forces that are typically located near the cell body. Cell forces are then summed over a user-defined sphere around the cell (12, 19). One advantage of this approach is that the exact position of the cell surface does not need to be known. A disadvantage is that balancing forces from sources outside the image stack can occur throughout the reconstructed force field.

In our new Python-based implementation (Saenopy), we regularize the reconstructed forces only within the reconstruction volume (which can be of the same size as the image stack, or larger) but not at its boundaries. This ensures that balancing forces from sources outside the image stack appear only at the surface of the reconstruction volume. Therefore, we do not need to limit the reconstructed cell forces to be within a defined distance around the cell (12, 19) (Fig. 1a middle) and instead can integrate them over the entire reconstruction volume, excluding the surface. This allows the size of the imaged volume to be greatly reduced without losing accuracy (Fig. 1c). It is also no longer required that force-induced matrix deformations near the image stack boundary must be negligible. Further, we image the cells for a sufficiently long time so that the cell has either migrated outside the image stack, or that the cell forces have changed their direction and magnitude multiple times so that the time average (median) of the matrix deformations corresponds to the undeformed state (Fig. 1b bottom).

From simulations of force-generating dipoles or quadrupoles with cell-like dimensions (15–30 µm) and forces, we verify the linearity of our method over a large force range (2 nN– 400 nN) even for highly non-linear matrices such as collagen (Fig. 1d), and confirm high sensitivity even in the presence of considerable imaging noise. For example, for a simulated cell that generates 20 nN (dipole) or 40 nN (quadrupole) of force, the typical displacement noise level of a confocal microscope of around 100 nm (*σ*_*x*_=41 nm, *σ*_*y*_=42 nm, *σ*_*z*_=99 nm) results in relative errors of contractility below 5% and relative errors of strain energy below 15%. (Fig. 1e).

The ability to obtain the force-free reference state without killing or relaxing the cells, combined with the ability to recover the traction field from an incomplete deformation field, allows us to measure the dynamic force generation of natural killer cells (NK92 cell line) during migration in 1.2 mg/ml collagen gels (shear modulus in the linear range of 100 Pa (SI Fig. 1–3), average pore size 4.4 µm (SI Fig. 4)). We record one image stack (123×123×123 µm, voxel-size 0.24×0.24×1 µm, acquisition time of 10 seconds using a resonance scanner (8000 Hz) and a galvo stage) every minute for 23 min. From these 23 image stack pairs, we compute the differential deformations using particle image velocimetry (PIV) (20), integrate them over time, and estimate the force-free matrix configuration (undeformed reference state) from the median deformation (Fig. 1b). Cell forces at each time point are then computed from the current matrix deformations relative to the undeformed reference state. Thus, this method allows for dynamic deformation measurements of highly motile cells with rapidly fluctuating traction forces. From the cell positions and cell shapes, we also extract cell velocity and aspect ratio (Fig. 2a–c).

**Fig. 2.**
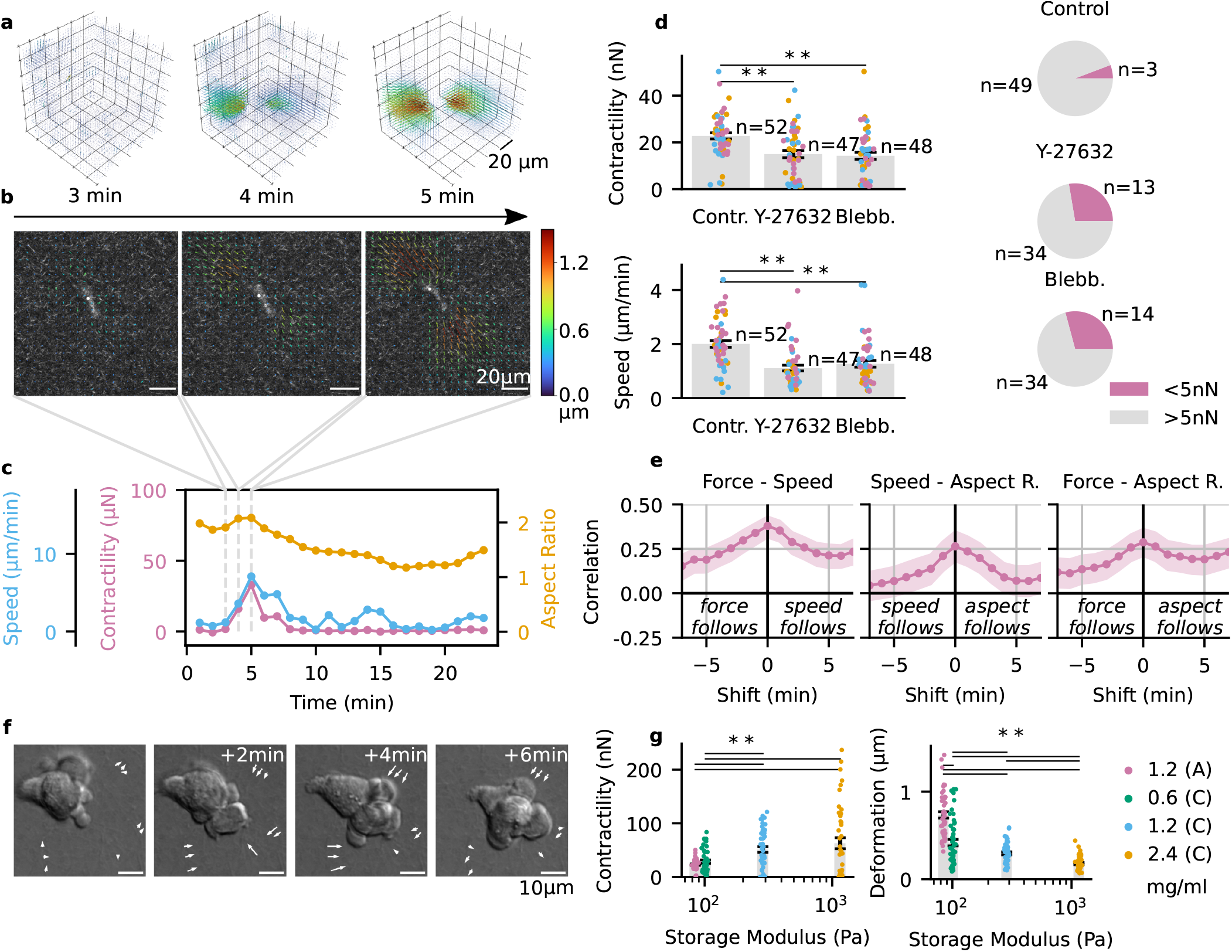
Contractile phases during natural killer cell migration. **a**, Matrix deformations (relative to the dynamic reference state) around a natural killer cell migrating in a collagen type I hydrogel at 3 time points selected from a 23 min time series. **b**, Corresponding maximum-intensity projected confocal reflection image overlaid with the projected maximum matrix deformation field. **c**, Cell speed (blue), cell aspect ratio (1: round, >1: elongated, orange) and cell contractility (magenta) over a 23 min period. Dashed gray lines indicate the time points shown in **a**,**b. d**, Cells treated with Y-27632 (10 μM) or Blebbistatin (3μM) show a decrease in maximum contractility (top left, mean±se), decrease in speed (bottom left, mean±se), and a decrease in the fraction of contractile cells (defined as contractility > 5 nN). Measurements of individual cells are shown as points, colors indicate independent experiments from different days. ** indicates p<0.01 for two-sided t-test. **e**, Cross-correlation between time courses of speed and force (left), speed and aspect ratio (middle), and aspect ratio and force (right). Signals were shifted relative to each other by ±5 min. The physical meaning of a positive correlation for different time points is indicated in the diagrams. Data points are mean values from 52 cells, shaded areas indicate one standard deviation determined by bootstrapping. **f**, Differential interference contrast (DIC) images show morphological changes of a NK cell during migration through collagen pores. Arrows indicate the local matrix deformation between consecutive frames. **g**, Cell contractility (maximum value within 23 min measurement period) and matrix deformation (99% percentile value of the absolute matrix deformation vectors is calculated per timestep. Then we choose the maximum value within 23 min measurement period) of NK92 cells embedded in collagen gels of different stiffness and collagen batches (A,C, see SI Fig. 1) (mean±se for 44-56 cells from three independent experiments; ** indicates p<0.01 for two-sided t-test, SI Fig. 6). The storage modulus of different collagen gels in the linear regime is derived from shear rheological measurements with 1% strain amplitude and and a frequency of 0.02 Hz (SI Fig. 3).

Around 78 % of NK cells that are elongated at t=0 migrate by more than one cell diameter (13 µm) during the observation time of 23 min (SI Fig. 5). When migrating, NK cells typically show an elongated morphology, whereas non-migrating cells remain round. The majority of migrating cells show bursts of substantial contractile forces between 5–50 nN (Fig. 2d, SI Video 20–26), lasting typically 3–5 min, followed by non-contractile phases (< 5 nN). Cells migrate both during contractile and non-contractile phases, but contractile phases are significantly (p < 0.01) correlated with higher speed and higher cell aspect ratio (Fig. 2c, e). Accordingly, speed and aspect ratio are also positively correlated (Fig. 2e). Myosin inhibition with blebbistatin and Rho-kinase inhibitor Y-27632 markedly reduce the magnitude and frequency of force bursts (Fig. 2d) and also reduce the migration speed (Fig. 2d).

Our discovery of substantial contractile forces in migrating immune cells that are approaching or even matching those of much larger cells, such as metastatic breast carcinoma cells (MDA-MB-231) or mouse embryonic fibroblasts (12, 22, 23), is surprising, as amoeboid migration is thought to be facilitated by shape changes and low frictional (traction) forces with the matrix (24–26). While our observation of long migratory phases with low tractions is in agreement with this notion, the occurrence of large traction force bursts points to a strong coupling of the cell’s contractile machinery to the extracellular matrix during these brief phases. Such strong coupling may be essential for migration through small matrix pores that pose a high steric hindrance and require large cell shape changes (Fig. 2f, SI Video 20–26).

Our finding of a strong coupling between the contractile machinery of immune cells and the mechanical properties of the extracellular matrix is supported by our finding of increasing tractions in collagen gels with increasing stiffness (Fig. 2g, SI Fig. 6). This behavior is reminiscent of mesenchymal cells that typically also respond with higher traction forces to higher matrix stiffness (18, 27, 28). In mesenchymal cells, this mechano-sensitive behavior is facilitated by integrin-mediated cell adhesions with the matrix. Our finding of a similar mechano-sensitive behavior in immune cells, together with our discovery of traction force bursts that are comparable in magnitude to those generated by mesenchymal cells, suggest that strong integrin-mediated adhesions may also be involved in immune cell migration through tight constrictions. This finding may have broad biological implications and adds to our current understanding of how immune cells migrate through tissue for their homing towards target sites in inflammation or cancer.

We next apply our method to measure traction forces of growth cones from retinal ganglion cell axons of African clawed frogs (*Xenopus laevis*). From 2D TFM measurements, it is known that forces of axonal growth cones are small (1–10 nN) (29–31) and highly dynamic (31, 32). In 3D collagen and PEG gels, axons growth cones have been reported to generate no measureable traction forces (33). Because axons are typically much larger than the 145 µm size of our image stacks, traction forces can be partially unbalanced. Despite these challenges, our method resolves traction forces of axonal growth cones in 3D of around ∼3 nN. As seen in immune cells, these forces act only intermittently, and longer periods of axon growth can occur without detectable traction forces (SI Fig. 7, SI Video 27).

In contrast to weakly contractile axon growth cones, fibroblasts are highly contractile and can induce extensive matrix deformations that play an important role in the progression of fibrosis or cancer (34–36). Even larger matrix deformations can be observed around spheroids or organoids consisting of hundreds or thousands of cells often. Spheroids and organoids are becoming increasingly important as sophisticated model systems for testing pharmacological compounds or to study biological processes during development, regeneration, or cancer progression (37–42).

To test our method in situations where traction forces are so large that they induce non-linear responses in the surrounding matrix, we measure primary liver fibroblasts (human hepatic stellate cells) as well as 100 µm sized multicellular mouse intestinal organoids (SI Fig. 7, SI Video 28, 29). In both cases, matrix deformations are large (∼10 µm) and reach beyond the size of the image stacks. Nonetheless, cell-generated forces (∼0.5–10 µN) are reconstructed with high sensitivity (Fig. 3) near the cell or organoid surface (SI Fig. 7). Due to the strong material non-linearity, large matrix deformations around spheroids and organoids can lead to slow convergence of the force reconstruction algorithm. Our new method takes advantage of the fact that the relative changes of the deformations and tractions between two consecutive time points are usually small. Therefore, we can use the deformation field from the preceding time point as the starting configuration for the force reconstruction of the subsequent time point. In the case of organoids, this approach speeds up the convergence of the algorithm by a factor of 5-50.

**Fig. 3.**
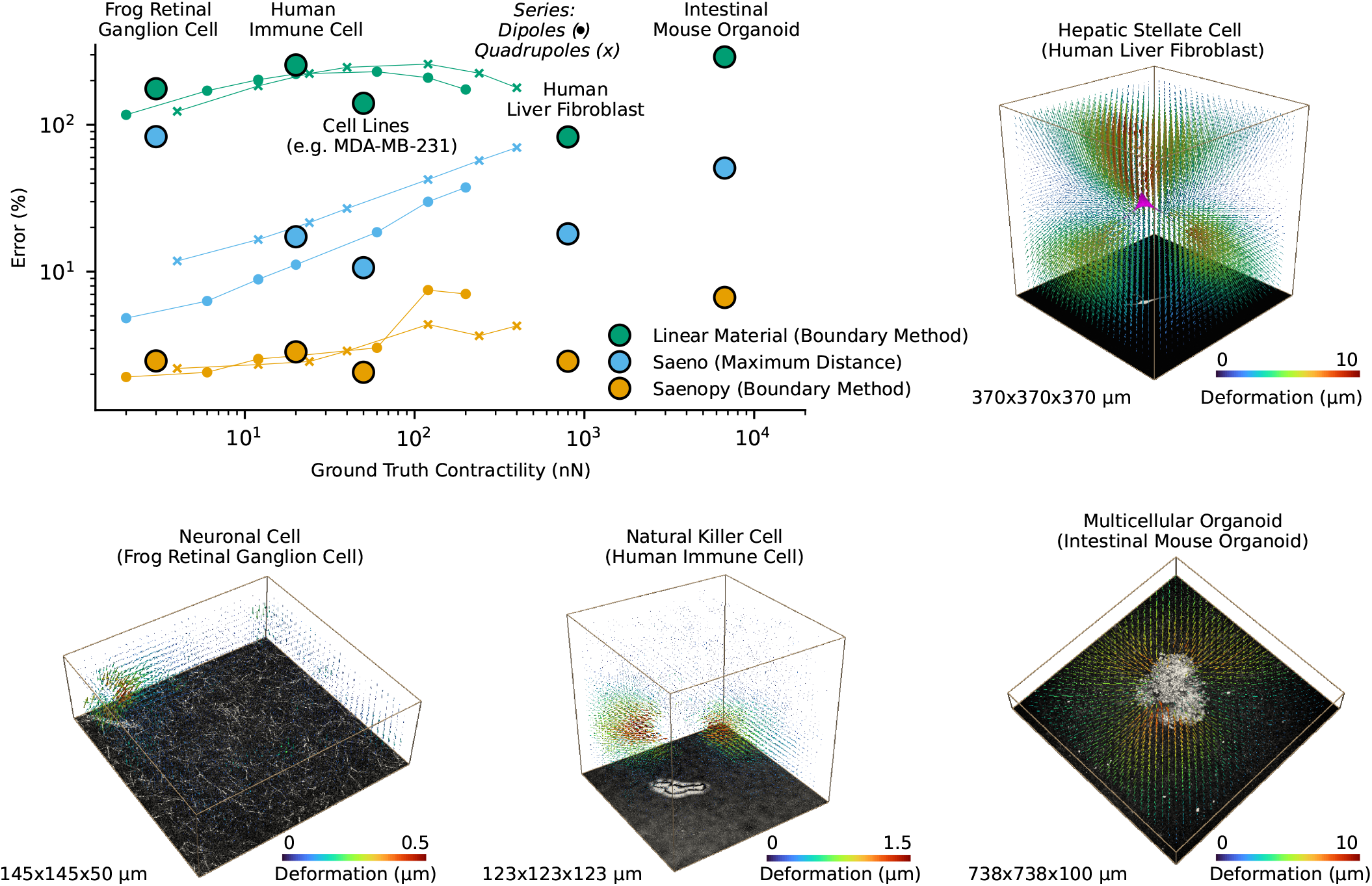
Accuracy and applicability of Saenopy. Relative error of traction force reconstruction for different cell types and ground-truth contractile forces. The forces are reconstructed from simulated matrix deformation fields based on monopoles, dipoles, and dodecahedra, with ground-truth forces, imaged volume, collagen rheology, and geometries taken from measurements of Xenopus retinal ganglion cell axon growth cones (monopols, SI Fig. 7, SI Video 27), NK92 cells (dipoles, Fig. 2, SI Fig. 7, SI Video 24-26), MDA-MB-231 breast carcinoma cells (dipoles, (12, 21)), primary human liver fibroblasts (dipoles, SI Fig. 7, SI Video 28), and primary intestinal mouse organoids (dodecahedra, SI Fig. 7, SI Video 29). The images show examples of the measured matrix deformation fields on which the simulations are based, with the corresponding confocal reflection images (axon growth cone, intestinal organoid), bright-field image (natural killer cell) or calcein-stained fluorescent image (hepatic stellate cell) shown below.

In summary, our method allows for the dynamic measurement of small to large 3D traction forces over prolonged time periods in linear or non-linear 3D fiber networks such as collagen, Matrigel, or fibrin. The method shows increased sensitivity and accuracy compared to previous 3D TFM methods (Fig. 3, SI Fig. 8-10), requires only a small confocal or bright-field image stack (SI Fig. 12, 11, SI Video 30), and is insensitive to forces from sources outside the imaged volume (Fig. 1c, SI Fig. 9). The method can be applied to single cells, cell colonies, spheroids, and organoids. For a user-friendly implementation of the method, we provide a Python-based open-source software platform (43) with a graphical user interface (SI Fig. 13,14).

## Methods

### Material model

Our method uses a previously developed material model for biopolymer hydrogels (12, 44). The material is filled with isotropically oriented fibers, each exhibiting the same non-linear stiffness (*ω*″) versus strain (λ) relationship according to Eq. (1) (Fig. 4). In particular, for small extensional strain in the linear range (0 < λ < λ_*s*_), fibers exhibit a constant stiffness *k*. For compressive strain (−1 < λ < 0), fibers buckle and show an exponential decay of the stiffness with a characteristic buckling coefficient *d*_0_. For larger extensional strain beyond the linear range (λ_*s*_ < λ), the fibers show strain stiffening with an exponential increase of stiffness with a characteristic stiffening coefficient *d*_*s*_.

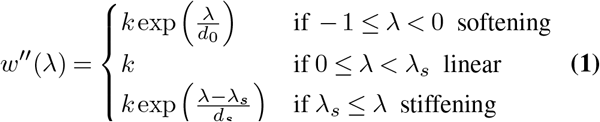

This description gives rise to a semi-affine behavior: Macroscopic material deformations cause non-affine deformations of the individual fibers, e.g. some fibers may buckle while others are stretched, depending on their orientation. Using a mean-field approach, we approximate the material behavior by averaging over many isotropically oriented fibers so that all spatial directions contribute equally.

**Fig. 4.**
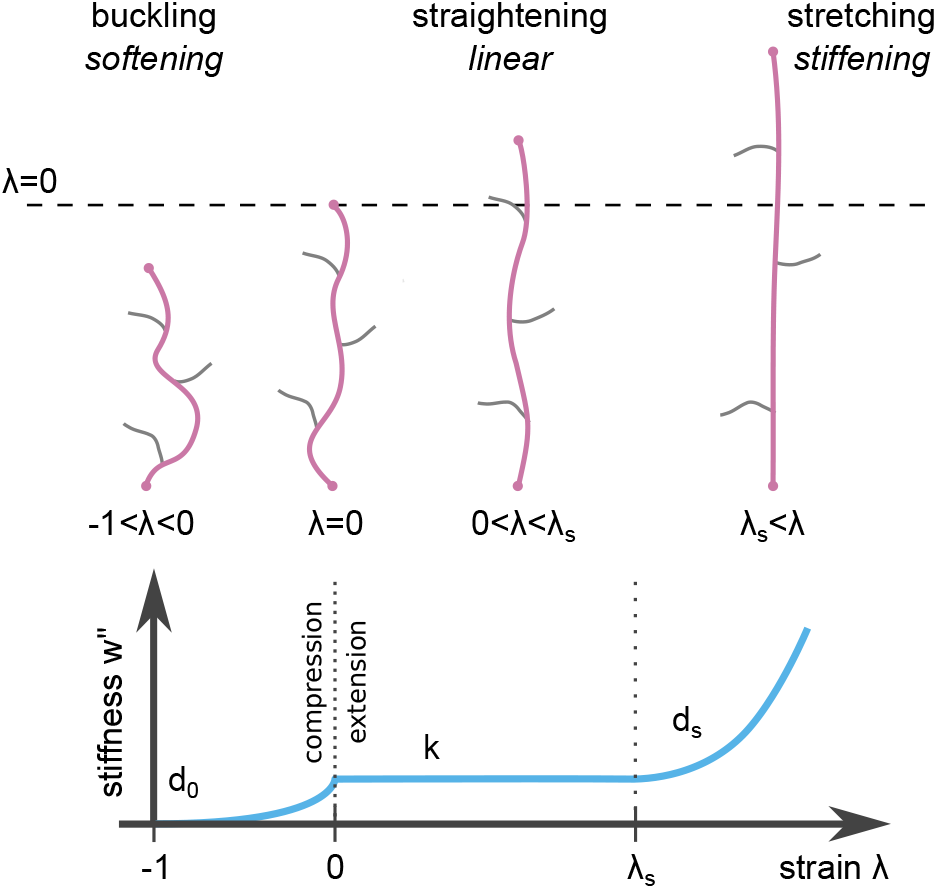
The nonlinear material model divides the mechanical response of individual fibers into a region where fiber stiffness (*ω*″) decreases exponentially with decreasing strain under compression (buckling), a region of constant fiber stiffness for small strains (straightening), and a region of exponentially increasing fiber stiffness for larger strains (stretching) (12).

Specifically, the energy function *w*(λ) is used to calculate the 239 energy density *W* stored in the material for a given deforma-240 tion gradient F, averaged over the full solid angle Ω (12, 44):

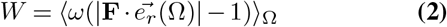

The strain λ of a unit vector 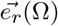 (representing a fiber), 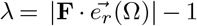, is used to calculate the energy *ω*(λ) stored in this spatial direction. The strain is averaged over the full solid angle Ω, approximated by a finite set (typically 300) of homogeneously and isotropically distributed unit vectors 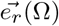.

### Finite element model

To describe the material behavior in the case of an inhomogenous deformation field, we tessellate the material volume into a mesh of linear tetrahedral elements *T*. The deformation field is then modeled as deformations 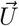 of the nodes of these elements. The total energy 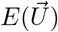 of the mesh is the sum over the energy density *W* of all tetrahedral elements *T* multiplied by their volume *V*_*T*_ :

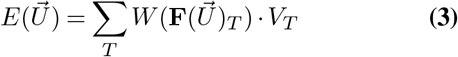

From the total energy 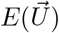, we calculate the stiffness matrix 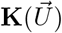 and the nodal forces 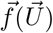 (12, 44):

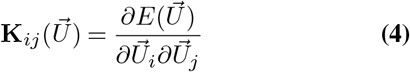

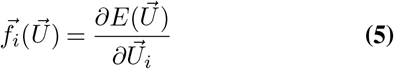

Index *i* represents the nodes of the finite element mesh, with index *j* representing the adjacent nodes.

### Mechanical characterization of biopolymer networks

We investigate the influence of three different collagen batches (A, B, C) and different collagen concentrations (0.6, 1.2 and 2.4 ml/ml) on the material behavior and the resulting material model parameters (SI Fig. 1). Two different rheological experiments are performed to measure the macrorheological behaviour of collagen hydrogels. First, we use a cone-plate rheometer (Discovery HR-3, TA Instruments, Milford, 20 mm cone-plate geometry, 2° angle, truncation gap 54 µm) to measure the stress-strain relationship of collagen gels for simple shear deformation. The collagen sample (85 µl) is polymerized inside the rheometer setup at 37°C for 30 min. A solvent trap filled with water is used to prevent evaporation and drying out of the sample. An initial frequency sweep (0.2–2 Hz) is performed at a low amplitude of 1%. Then, the stress-strain relationship is measured for a strain ramp from 0% to 100% with a constant strain rate of 1%/s.

In a second experiment, we measure the vertical contraction of collagen gels in response to horizontal uniaxial stretching. Stretching is performed using 2x2 cm flexible polydimethylsiloxane (PDMS) membranes (Sylgard184; crosslinker-to-base ratios of 1:14; cured for 24 h at 65°C, Sigma-Aldrich, St. Louis) as described in (45). The PDMS membrane is pre-stretched to 5% and coated with 0.5 mg/ml Sulfo-SANPAH (Thermo Fisher Scientific, Waltham) followed by 5 min UV treatment and 3x washing with phosphate buffered saline (Gibco PBS, Thermo Fisher Scientific, Waltham). Next, 700 µl of unpolymerized collagen solution is mixed with 4 µm silica beads (Kisker Biotech, Steinfurt) that serve as fiducial markers (5 µl beads per 1 ml). The collagen solution is transferred to the PDMS membrane, and after 20 min polymerization at 37°C and 5% CO_2_, another 5 µl of the silica bead suspension is pipetted onto the gel to mark the gel surface. The collagen is polymerized for a total of 1 hour, after which 1 ml of DMEM medium (1 g/l glucose, Thermo Fisher Scientific, Waltham) is added on top. The collagen gel is stretched uniaxially in 1% step increments using a custom made stepper motor-driven cell stretching device (46) mounted on an epifluorescence microscope (Leica DMI6000 CS equipped with a 20x Leica HCX PL Fluotator objective, Leica, Wetzlar). For each stretch increment, we measure the gel thickness by focusing on the top and bottom surface of the gel (highest and lowest layer of beads) and correct for the refractive index of water.

For collagen batch D (used only for axon growth cone measurements), the stress-strain relationship is determined using a rheometer equipped with a 25 mm diameter cone-plate geometry (MCR302e, AntonPaar, Graz). The polymerisation of 50 µl of hydrogel between the rheometer plates is monitored for 90 minutes at 20°C using an oscillatory strain of 0.5%. After polymerization, a frequency sweep (0.1–100 rad/s) with 0.5% strain is performed, followed by a strain ramp at a constant strain rate of 0.5%/s (SI Fig. 2).

Because a cone-plate rheometer or uniaxial stretching device may not be available in every laboratory, and hence the correct material parameter for a given collagen batch may be unknown, we also quantify the effects of using erroneous material parameters on cellular force reconstruction. (SI Fig. 16-17). Furthermore, we explore the possibility to derive the linear stiffness parameter k_0_ of the material model, which most sensitively affects force reconstruction, from oscillatory cone-plate rheology measurements in the linear range (SI Fig. 3).

### Estimating material parameters from rheological measurements

As the material model is based on the mechanical behavior of microscopic biopolymer fibers, the model parameters cannot be directly extracted from macroscopic rheological measurements. Instead, we simulate the macrorheological experiments using different model parameters until a best-fit is achieved (SI Fig. 1). Data from uniaxial stretching are required to fit the buckling coefficient *d*_0_ of the material. For the other parameters, either data from a shear rheometery or an extensional rheometry experiment are sufficient as long as the strain range exceeds the linear range of the material (43).

### Collagen gel preparation

Collagen type 1 hydrogels are prepared from acid-dissolved rat tail (R) and bovine skin (G1) collagen (Matrix Bioscience, Mörlenbach). Both collagen types are mixed ice-cold at a mass ratio of 1:2 and are dissolved in a dilution medium containing 1 vol part NaHCO_3_, 1 vol part 10× DMEM and 8 vol parts H_2_O, adjusted to pH 9 using NaOH. The final collagen concentrations are adjusted to 0.6 mg/ml, 1.2 mg/ml or 2.4 mg/ml using dilution medium. The the final mixture is then adjusted to pH 9 using NaOH. Collagen gels are polymerized for 1 h at 5% CO_2_ and 37°C. For axon growth cone measurements, collagen type 1 hydrogels are prepared from acid-dissolved bovine collagen (TeloCol-10, Advanced Biomatrix, Carlsbad). 6 parts Leibovitz L-15 medium (Gibco, Thermo Fisher Scientific, Waltham), 3 parts autoclaved double-distilled water, and 1 part collagen solution are prepared for a final collagen concentration of 1 mg/ml. HEPES buffer (15 mM, Sigma-Aldrich, St. Louis), 100 units/ml penicillin-streptomycin and 0.25 µg/ml amphotericin B (Lonza, Basel) are added, and the final mixture is adjusted to pH 7.3 using NaOH. Collagen gels are polymerized for 1 h at ambient CO_2_ and 20°C.

### Cell culture and 3D traction force experiments. Immune cells

NK92 cells (ATCC CRL-2407) are cultured at 37 °C and 5% CO_2_ in Alpha-MEM medium (Stem-cell Technologies) with 15% fetal calf serum, 15% horse serum, 500 IU/ml human IL2-cytokine and 1% penicillin-streptomycin solution (10.000 Units/ml penicillin, 10.000 µg/ml streptomcycin). Cells are suspended in 3 ml ice-cold collagen (1.2 mg/ml) solution (66.000 cells/ml), transferred to a 35 mm Nunc dish (Thermo Fisher Scientific, Waltham) and polymerized at 37°C, 5% CO_2_ for 1 h. After polymerization, 2 ml of pre-warmed medium are added, and the sample is transferred to an upright laser-scanning confocal microscope (Leica SP5, equipped with an environmental chamber to measure at 37°C, 5% CO_2_). To achieve high time resolution, we use a resonance scanner (8000 Hz) in combination with a galvo-stage. Image stacks (123×123×123 µm with a voxel-size of 0.24 ×0.24×1 µm) around individual cells are recorded in bright-field and confocal reflection mode using a 20x water dip-in objective (HCX APO L 20x/1.00 W, Leica, Wetzlar). The acquisition time for an individual image stack is below 10 seconds. Up to four individual cells at different positions are recorded every minute over a period of 23 min.

### Liver fibroblasts

Human hepatic stellate cells (HUCLS, Lonza, Basel) are cultured at 37°C, 95% humidity and 5% CO_2_ in human hepatic stellate cell growth medium (MCST250, Lonza, Basel). 3 ml collagen solution (1.2 mg/ml, Batch C) are mixed with 35 000 cells and transferred to a 35 mm Petri dish (Greiner AG, Austria). Samples are polymerized at 37°C for 60 minutes, after which 2 ml of cell culture medium are added. Two days after seeding, the medium is replaced with fresh medium, and cells are stained using 2 µM calcein AM (Thermo Fisher Scientific, Waltham). Samples are transferred to an upright laser-scanning confocal microscope (Leica SP5, Wetzlar) equipped with a 20x water dip-in objective (HCX APO L 20x/1.00 W, Leica, Wetzlar) and an environmental chamber (37°C with 5% CO_2_). Cubic image stacks (370 µm)^3^ with a voxel-size of 0.72×0.72×0.99 µm are recorded around the cells in bright-field, fluorescence and confocal reflection mode. The force-free reference state is acquired ∼ 30 min after drug-induced relaxation of cell forces using 10 µM cytochalsin D (Sigma-Aldrich, St. Louis). Force reconstruction is performed on confocal reflection image stacks using an element size of 14 µm for performing particle image velocimentry (PIV), a PIV window-size of 35 µm, and a signal-to-noise filter of 1.3. The finite element model has an element size of 14 µm. The regularization parameter α is set to 10^10^, and the material properties for collagen Batch C are used (SI Fig. 1).

### Axon growth cones

Xenopus medium is prepared from 6 parts Leibovitz L-15 medium and 4 parts autoclaved double-distilled water, 100 units/ml penicillin-streptomycin, and 0.25 µg/ml amphotericin B (Lonza, Basel). Primary eye primordia explants are dissected from Xenopus laevis embryos as previously described (47). In brief, Xenopus embryos develop from in vitro fertilized eggs at ambient CO_2_ and 20°C in 0.1×Marc’s Modified Ringer’s Solution (MMR). After typically 6 days of incubation, embryos at a development stage of 35/36 (48) are transferred to a polydimethylsiloxane (PDMS)-coated dissection dish (Sylgard 184, Dow, Midland) filled with equal parts of MS-222 anaesthetization medium (Sigma-Aldrich, St. Louis) and Xenopus medium.

Explants are dissected under a preparation microscope (Leica, Wetzlar) under circular movements of dissection pins around the eye to remove the surrounding skin, and shear movements to extract the retinal explants. Dissected eyes are stored for no longer than 1 hour in Xenopus medium. The retinal explants are suspended in 50 µl Xenopus medium, and 1 ml of ice-cold collagen 1.0 mg/ml solution (Batch D) is prepared and mixed gently with the explant-medium solution. To prevent the explants from sinking to the bottom, the mixture is pre-polymerised for approximately 30 seconds at room temperature before it is transferred to a 35 mm Petri dish (Ibidi, Gräfelfing). The sample is polymerized for 2.5 h at room temperature, after which 2 ml of Xenopus medium is added and the sample is stored at ambient CO_2_ and 20°C. After 29 hours, the sample is transferred to a laser-scanning confocal microscope (TCS SP8, Leica, Wetzlar) equipped with an oil-immersion/ objective (HC PL APO 40x/1.30, Leica, Wetzlar). Image stacks (145×145×50 µm with a voxel-size of 0.14×0.14×1 µm) around a developing axon growth cone are recorded in bright-field and confocal reflection mode every 5 min for a duration of 2 h. Xenopus experiments are conducted in accordance with the UK Animals (Scientific Procedures) Act 1986.

Force reconstruction is performed on confocal reflection image stacks (SI Video 27) using a PIV element size of 4.8 µm, a PIV window-size of 12 µm and a signal-to-noise filter of 1.3. The finite element model has an element size of 4.8 µm. The regularization parameter α is set to 10^11^, and the material properties for collagen batch D are used (SI Fig. 2).

### Intestinal organoids

Experiments involving mice have been approved by the Institutional Animal Care and Use Committee of the University of Erlangen-Nuremberg and by the Government of Mittelfranken (Würzburg, Germany). Small intestines are collected from C57BL/6J mouse (Strain #:000664, The Jackson Laboratory), washed with iced-cold PBS, and longitudinally opened. Villi are mechanically removed with the help of a glass coverslide, the tissue is then cut into 1 cm pieces and incubated in chelation buffer (2 mM EDTA in PBS) with gentle shaking in a rocking rotator for one hour. After several washing steps with iced-cold PBS, the supernatant is filtered through a 70 µm cell strainer to collect isolated crypts. They are embedded in basement membrane extract hydrogels (Cultrex RGF, Biotechne, Minneapolis) and placed in a pre-warmed 48-well plate (Nunc delta surface, Thermo Fisher Scientific, Waltham). The organoids are cultured in Advanced DMEM/F12 medium (Gibco, Thermo Fisher Scientific, Waltham) supplemented with N-Acetylcistein (1 mM, Sigma-Aldrich, St. Louis), mEGF (50ng/ml, Immuno tools, Friesoythe), B27 and N2 supplement (1x, Life Technologies, Carlsbad), Primocin (50µg/ml, InvivoGen, Toulouse), and conditioned medium from R-spondin1 and mNoggin producing cells (batch dilution is defined experimetally based on titration using organoid survival as read-out). The medium is changed every 3 days, and organoids are passaged every 7–10 days. For harvesting and passaging, organoids are dissociated and seperated from the surrounding matrix by repeated pipetting and centrifugation (300g at 4°C for 5 min).

For traction force measurements, organoids are embedded in 1.2 mg/ml collagen type I hydrogels (Batch C). A two-layered gel is used to prevent sinking of organoids to the bottom during polymerization (21). In brief, 1.5 ml collagen solution are transferred to a 35 mm dish and polymerized for 20 minutes (5% CO_2_, 37°C). The supernatant medium from harvested organoids is removed, and organoids are diluted in another 1.5 ml of collagen solution. The mixture is then added on the pre-polymerized collagen layer. After 1 hour of polymerization, 2 ml of medium are added. Prior to imaging, organoids are stained with 2 µM calcein AM (Thermo Fisher Scientific, Waltham) and transferred to an upright laser-scanning confocal microscope (Leica SP5, 37°C, 5% CO_2_). Image stacks (738×738×100 µm with a voxel-size of 0.74×0.74×2 µm) around an organoid are recorded every 20 minutes in bright-field, fluorescence and confocal reflection mode using a water dip-in objective (HCX APO L 20x/1.00 W, Leica, Wetzlar). Organoid-generated forces are relaxed after ∼ 24 hours using 10 µM cytochalsin D (Sigma-Aldrich, St. Louis) and 0.1% triton x-100 (Sigma-Aldrich, St. Louis). The force-free reference state is acquired ∼ 2 hours later (SI Video 29).

Force reconstruction is performed on confocal reflection image stacks using a PIV element size of 30 µm, a PIV window-size of 40 µm and a Signal-to-noise filter of 1.3. The finite element model has dimensions of (738 µm)^3^ with an element size of 30 µm. Regularization parameter α is set to 10^10^, and material properties of collagen Batch C are used (SI Fig. 1).

### Bright field microscopy

During migration through the collagen network, NK92 cells are imaged in differential interference contrast mode (Leica DMI6000B, Wetzlar, with an HC PL APO 63x/1,40 oil immersion objective.) with 2 seconds between consecutive images (Fig. 2f).

### Measurement of matrix deformations

Cell-induced matrix deformations are measured from confocal reflection image stacks of the collagen network using the open-source particle image velocimetry algorithm OpenPIV (20) that we developed specifically for analyzing 3D image stacks. Alternatively, matrix deformation can also be measured from bright-filed image stacks of collagen gels either containing embedded microbeads (SI Fig. 11), or no beads but with sufficient image contrast from the individual collagen fibers (SI Fig. 12, SI Video 30). Saenopy offers two options for quantifying matrix deformations. First, matrix deformations can be calculated between a deformed state and a force-free reference state (Fig. 1b top). The reference image stack is typically recorded after drug-induced force relaxation using e.g. high concentrations of cytochalasin D (12, 19, 21). For fast moving cells, Saenopy offers a second option for obtaining the reference state from a sufficiently long time series of image stacks (Fig. 1b bottom): First, the change in matrix deformations between consecutive image stacks, which correspond to the time derivative of the matrix deformations, are calculated at each voxel position using OpenPIV (20). We then calculate the cumulative sum of the time derivative at each voxel, which corresponds to the absolute matrix deformation apart from an offset. To remove the offset, we subtract the median deformation at each voxel position, assuming that the median matrix deformations around a fast moving cell tend towards zero, when observed over a prolonged time period. For the NK92 cells in this study, we employ the second option. By comparing a region of the collagen matrix before and after an NK cell has passed through, we confirm that these cells do not permanently remodel the matrix.

For the OpenPIV algorithm, we use a window-size of 12 µm with an overlap of 60% (corresponding to an element size of 4.8 µm). Deformation vectors with low confidence (defined by OpenPIV for a ratio < 1.3 between the global maximum of the image cross correlation divided by the second-highest local maximum value) are ignored. The accuracy with which we can measure matrix deformations with our setup is estimated by imaging collagen gels without cells. We record image stacks at 28 non-overlapping positions in the gel (each 123×123×123 µm), each over a period of 23 min with a time interval of 1 min and a voxel-size of 0.24 × 0.24×1 µm as in the cell experiments. We then compute the deformation field at every voxel position as described above, and the accuracy (noise level) is calculated as the standard deviation σ of all deformation values, separately evaluated in x,y, and z-direction. We obtain values of σ_*x*_=41 nm, σ_*y*_=42 nm, σ_*z*_=99 nm.

### Cell tracking

Cell shape and position at each timepoint are extracted from intensity-projected bright-field image stacks. We compute the local entropy of the band-pass filtered images, subtract the background using regional maxima filtering, binarize the image using Otsu-thresholding (49), and apply binary morphological operations to remove small objects and holes. From the binarized object (which is the segmented cell), we calculate the area, the x,y position (center of mass), and the aspect ratio defined here as the major axis divided by the minor axis of an ellipse fitted to the cell shape. If a cell reaches the edges of the image, this cell is not tracked further.

### Correlation analysis

We cross-correlate the time development of cellular contractility, cell speed, and cell aspect ratio for all NK92 cells under control conditions (Fig. 2d,f). Since cell velocity and cell forces are calculated from the difference in cell position and matrix deformations between two consecutive image stacks, whereas the aspect ratio is obtained for each single image stack, we linearly interpolate the aspect ratio between two consecutive image stacks before cross-correlation. We then calculate the Spearman rank correlation coefficient between speed and contractility, between speed and aspect ratio, and between contractility and aspect ratio, for time shifts from -7 to +7 min. The error intervals are determined using bootstrapping with a sample-size of 1000. For computing cross-correlations of speed and aspect ratio with the contractility signal, only time points with positive contractility are considered.

### Force reconstruction

To solve the inverse problem of reconstructing cellular forces 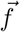 from the measured 3D deformation field 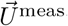 (Eq. 5), we use an iterative approach that varies the simulated matrix deformation field 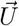 with the aim to minimize a target function 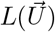. This target function is computed from the difference between the simulated matrix deformations 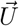 and the measured matrix deformations, 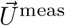, plus a Thikonov regularization term 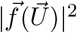 (12, 50, 51) to constrain the force field 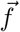:

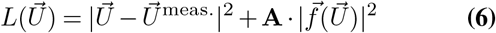

The force field 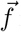 is computed from the simulated deformation field 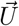 using Eq. 5. The diagonal regularization factor matrix **A** has weights that favor large forces (presumably generated by the cell) and penalizes small forces (presumably caused by measurement noise) (12, 52) (Eq. 7). At the boundary (surface of the simulated volume), we set the regularization factor to zero (see Section Boundary forces). The index i represents the nodes of the finite element mesh.

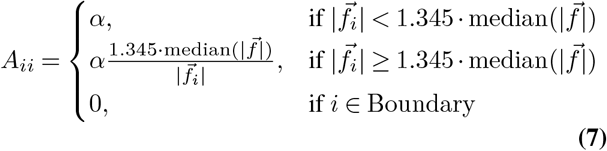

As 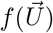 is non-linear, 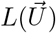 cannot be minimized easily. We apply a first-order Taylor series expansion of the displacement 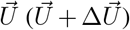, using the stiffness matrix **K** (Eq. 4) (44, 51).

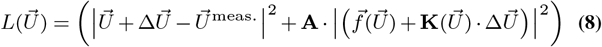

The 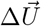 that minimizes this equation satisfies the following normal equation (51).

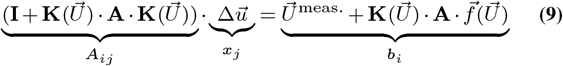

This linear equation (of the form A_*ij*_·x_*j*_ = b_*i*_) is solved using the conjugate gradient method to obtain a value for 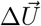. Because of the pronounced non-linearity of the problem, for the next iteration cycle we update the simulated deformation field only by a fraction (stepper) of 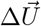 (typically 0.33):

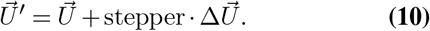

With the new displacement 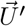, the stiffness matrix 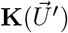 (Eq. 4), the nodal forces 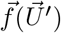 (Eq. 5), and the weight matrix 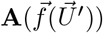 (Eq. 7) are updated, and the linear Taylor expansion (Eq. 9) is solved again. This procedure is iterated until the total energy 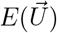 (Eq. 3) of the system converges. Convergence is reached when the standard deviation of 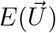 divided by the mean of 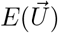 (coefficient of variation) for the last 6 iterations is below a convergence criterion τ (typically 0.01).

From the resulting force vectors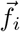 at all nodes i, we compute the coordinates of the force epicenter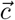. To find it, we minimize the magnitude Q of the cross product of the force field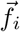 with the vectors from the nodes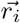 pointing to the epicenter coordinates 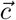 (12, 44):

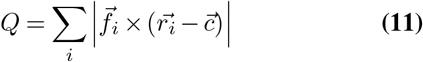

We determine the cellular contractility as the sum of the projected forces in the direction of the force epicenter (12, 44):

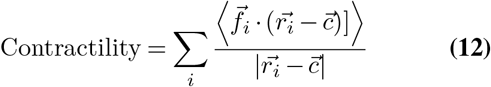

In this study, we perform the force reconstruction of migrating NK92 cells using the regularization parameter α = 10^10^ (Eq. 7), which maximizes contractility and is stable against noise both for the experimental setup and for simulated cells (SI Fig. 18). We interpolate the 3D matrix deformations onto a cubic grid with a grid-size of 4 µm (in the case of single cells, or larger as specified in Methods), and tesselate cubes into 6 tetrahedra (SI Fig. 19).

### Boundary forces

Because the boundary of the simulated, finite-sized volume is fixed, the reconstructed force field contains cell-generated forces from the bulk, as well as distributed counter-forces from the surrounding matrix that appear at the volume boundary. The individual force vectors of the counter-forces are small compared to cell forces. Hence, the small-force penalization scheme (Eq. 7, first and second case) would bundle them into fewer, larger force vectors located in the bulk where they could interfere with cell forces. To avoid this, we set the regularization weights *A*_*ii*_ to zero for all tetrahedral mesh nodes at the boundary (surface) of the simulated volume (Eq. 7, third case). Hence, we exclude the surface of the mesh from the regularization cost function so that virtually all counter-forces only appear at the volume boundary and not in the bulk (Fig. 1a). Saenopy can therefore directly sum all the forces in the bulk of the mesh as they are contractile cell forces, without the need to arbitrarily define a maximum radius around the cell center for the summation of forces. This improves robustness and accuracy especially for small stack sizes (Fig. 1c,e).

### Simulations of force-generating dipoles and quadrupoles

We simulate the 3D deformation fields around force-generating dipoles and quadrupoles in a 1.2 mg/ml collagen matrix (Batch A; SI Fig. 1) with the iterative scheme described above, with the following modifications: The displacements at the mesh boundary are set to zero, the forces at the dipole or quadrupole points are given, and the forces in the bulk are set to zero. To simulate contractile dipoles, we add two opposing point forces 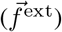with a distance of 15 µm or 30 µm. To simulate contractile quadrupoles, we add 4 inward pointing forces 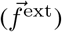at the nodes of a regular tetrahedron with an edge length of 15 µm or 30 µm. We now iteratively vary the deformation field until the simulated forces match the imposed force field using the conjugate gradient method as described above. However, instead of the Eq. 9, we solve the following equation:

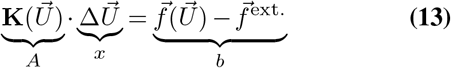

and update the deformation field 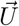 after each iteration according to Eq. 10, as well as the the stiffness matrix 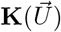 (Eq. 4) and the nodal forces 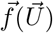 (Eq. 5), until the total Energy 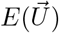convergences (12, 44).

Point forces are applied to the nearest possible nodes for a grid-size of 4 µm and a stack volume of (150 µm)^3^. We simulate various contractilities (2 nN–400 nN) with or without added Gaussian noise with a defined standard deviation σ to the deformation field.

To benchmark the accuracy of different 3D TFM reconstruction methods (Fig. 3), we simulate the collagen deformations around artifical cells according to Eq. 13 and then perform force reconstruction. Axon growth cones are simulated by monopoles, immune cells, hepatic stellate cells and cancer cells by dipoles, and intestinal organoids by dodecahedrons, each with corresponding contractilities, spatial dimensions and material parameters taken from experiments. For force reconstruction, we use standard Saenopy (Boundary Method), Saenopy in combination with a purely linear elastic material model (= linear method), and Saeno (Maximum Distance method, whereby *r*_*max*_ is optimized for the best possible agreement of simulated and reconstructed contractility values).

## Limitations

### Resolution/Heterogeneities

The spatial resolution of the recovered cellular forces is limited by the pores size of the matrix, which in turn limits the window size of the particle image velocimetry algorithm and the size of the finite elements. Moreover, the fibrous structure of the matrix naturally introduces local heterogeneities at length scales even above the pore size. Therefore, the spatial resolution is on the order of 5-10 times the average pore size, i.e. around 30 µm in collagen gels (12). However, deformations in collagen are propagated over long distances within the matrix so that the global contractility of the cell can still be accurately recovered.

### Viscoelasticity/Plasticity

The material model of our method accounts for non-linear elastic matrix deformations, but ignores viscoelastic or plastic properties. This is justified for moderate strains (below 20%) where collagen behaves predominantly as an elastic material (SI Fig. 3). Moreover, after drug-induced relaxation of cellular forces, the recoil of the matrix deformations are driven by the release of elastically stored strain energy so that an elastic material model is appropriate. Potential problems arise only if the force-free matrix configuration is calculated from images taken at the beginning of the experiment (or from the median deformations) so that non-elastic matrix deformations, if present, can contribute to the total deformations. In this case, a purely elastic model overestimates the forces. For highly motile immune cells, however, we do not observe permanent deformations of the collagen matrix after the cell has migrated through it.

## Data Availability

The software (Saenopy) and all dependent packages are are available on GitHub as an open-source python package with a graphical user interface (43). Figures are created using the python package Pylustrator (53) and PyVista (54). The data of this study are available upon request from the corresponding author. Rheological measurement of 1.2 mg/ml collagen gels of Batch A were previously published in (55).

## Author contribution

Methodology: RG, DB, MC, BF, CM. Software: RG, DB, MC, AB. Rheology: DB, MC, LB, SB, NG. Immune cell experiments: LB, TC, DB, CV. Organoid experiments: PAN, DB, RLP. Neuronal cell experiments: KF, NG. Data Analysis: DB, LB, CM, RG. Writing: DB, BF, MC, RG.

## ACKNOWLEDGEMENTS

This work was funded by the German Research Foundation (DFG; project 326998133 – TRR-SFB 225 – subprojects A01, B09 and C02; project 375876048– TRR-SFB 241 – subprojects A07 and C04; project 461063481 – LO 2465/6-1; project 414058251 – SPP-1782, LO 2465/2-1), the National Institutes of Health (HL120839), the Emerging Fields Initiative of the University of Erlangen–Nuremberg, and the Alexander von Humboldt Foundation (Humboldt Professorship, K.F.). We thank the ENB Biological Physics program of the University Bayreuth for support, Ricardo Henriques for the latex layout, Ingo Thievessen for help with DIC imaging, and Ronny Reimann for designing the Saenopy logo.

---

**Supplementary Information 1:**
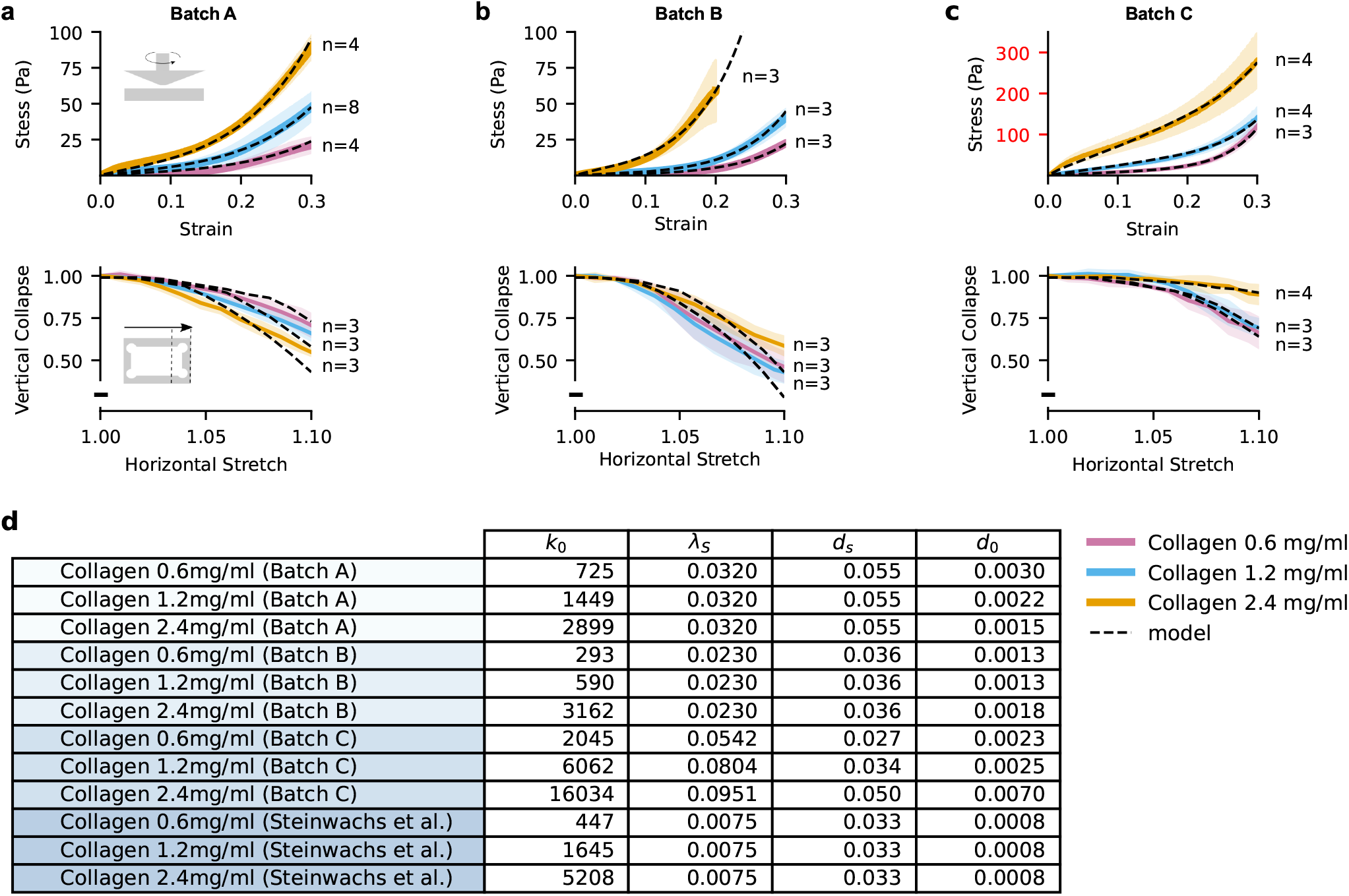
Rheology and material model for collagen Batch A, B, C. Collagen hydrogels from three different batches (**a, b, c**) at different concentrations of 0.6 mg/ml (purple), 1.2 mg/ml (blue), and 2.4 mg/ml (orange) are characterized using cone-plate shear-rheological measurements (top) and uniaxial stretch-experiments (bottom) as described in the Methods section of the main text. Solid lines represent the mean, shaded areas represent one standard deviation. Dashed lines show the fit of the finite element model to the data. **d**, Model parameters are extracted from the fit for three different batches (Batch A-C). Additionally, previously measured parameters from Steinwachs et al. (12) are given for comparison. For each batch, we performed both individual fits to the data for each concentration separately, or global fits where the parameters *λ*_*s*_, *d*_*s*_, and *d*_*0*_ were the same for all or some of the concentrations. Global fit parameters were preferred if the fit quality was comparable to individual fits, in order to reduce the number of free fit parameters.

**Supplementary Information 2:**
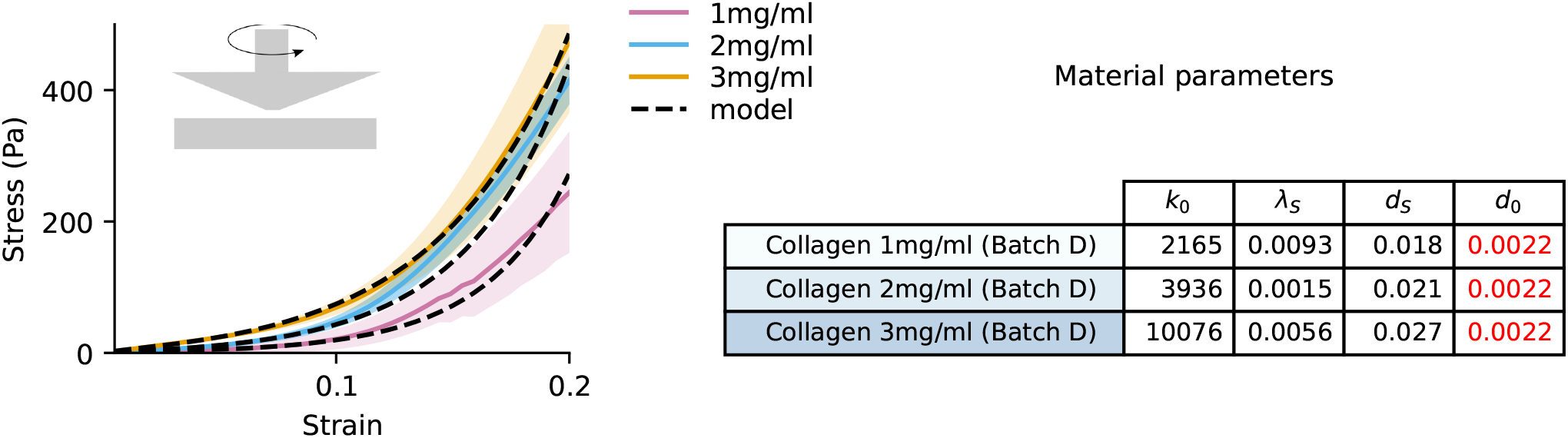
Rheology and material model for collagen Batch D. Collagen hydrogels from Batch D (used only for axon growth cone experiments) are characterized at different concentrations of 1 mg/ml (purple), 2 mg/ml (blue), and 3 mg/ml (orange) using cone-plate shear-rheological measurements as described in the Methods section. Solid lines represent the mean value, shaded areas represent one standard deviation (n = 3). Dashed lines show the fit of the finite element model to the data. The table lists the material model parameters for each concentration. The buckling parameter *d*_*0*_ is selected based on 1.2mg/ml collagen Batch A (red values, see SI Fig. 1).

**Supplementary Information 3:**
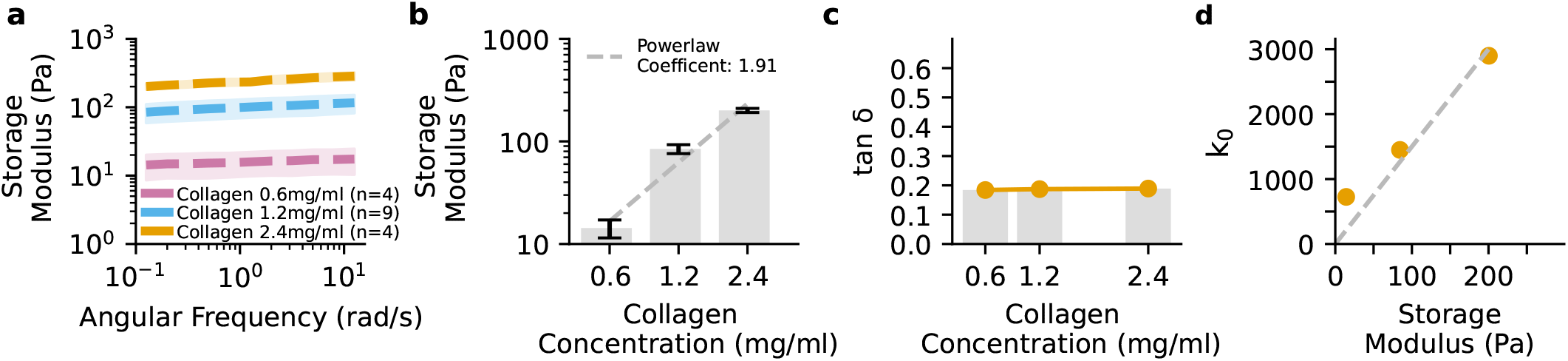
Small amplitude rheology. Storage modulus derived from frequency sweeps with a cone-plate rheometer at 1% strain amplitude for different collagen concentrations (Batch A, see SI Fig. 1). Dashed lines indicate mean values, and shaded area indicate ± one standard deviation. Storage modulus (mean value at 0.02 Hz) scales with collagen concentration according to a power-law with exponent of 1.91 in agreement to previously predicted and measured values (56**?**). Bars indicate mean±se and dashed line indicates powerlaw fit curve. **c**, Loss tangent *δ* (loss modulus *G″* divided by storage modulus *G′*, averaged from 0.02-2 Hz) remains below 0.2 for all collagen concentrations, indicating predominantly elastic behavior. **d**, The FE model parameter *k*_*0*_ (indicating the linear stiffness of the collagen fibers) for different collagen concentrations (see SI Fig. 1) increases approximately linearly with the storage modulus *G′* of the collagen gels (measured at 0.02 Hz at a strain amplitude of 1%). The gray dashed line indicates the predication from continuum mechanics, where *k*_0_ = *6E*, with YounG′s modulus *E = G* ·2· (1 + *v)* and Poisson ratio *v =* 0.25 for linear elastic, isotropic fiber networks (12). Hence, *k*_0_ = 15 · *G*.

**Supplementary Information 4:**
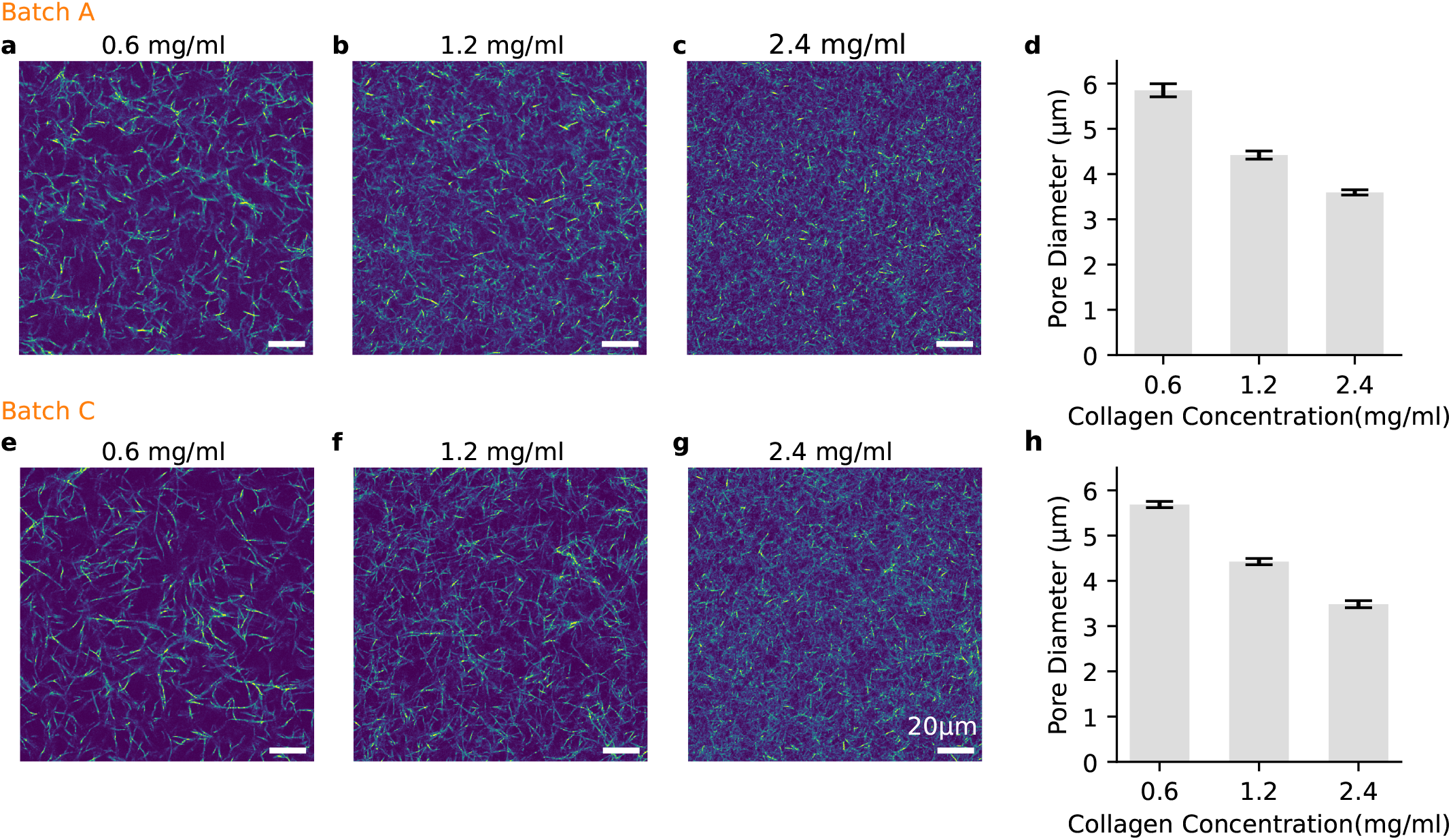
Microstructure of collagen networks. The collagen fiber structure of two different collagen batches (**a**-**c**: Batch A; **e**-**g**: Batch C) is imaged using confocal reflection microscopy for three different collagen concentrations (imaged volume of 160 ×160 ×200 μm with voxel-sizes of 0.314×0.314×0.642 μm). Images show a single slice. 3D pore diameters are computed from the covering radius transform as described in (57, 58). The mean pore diameters for each collagen concentration (**d**,**h**) are calculated from 8 different regions within an stack (80x80x100 μm with 0.314x0.314x0.642 μm voxel-size). The error bars represent the standard deviation of the mean pore diameter between different regions of the imaged stack.

**Supplementary Information 5:**
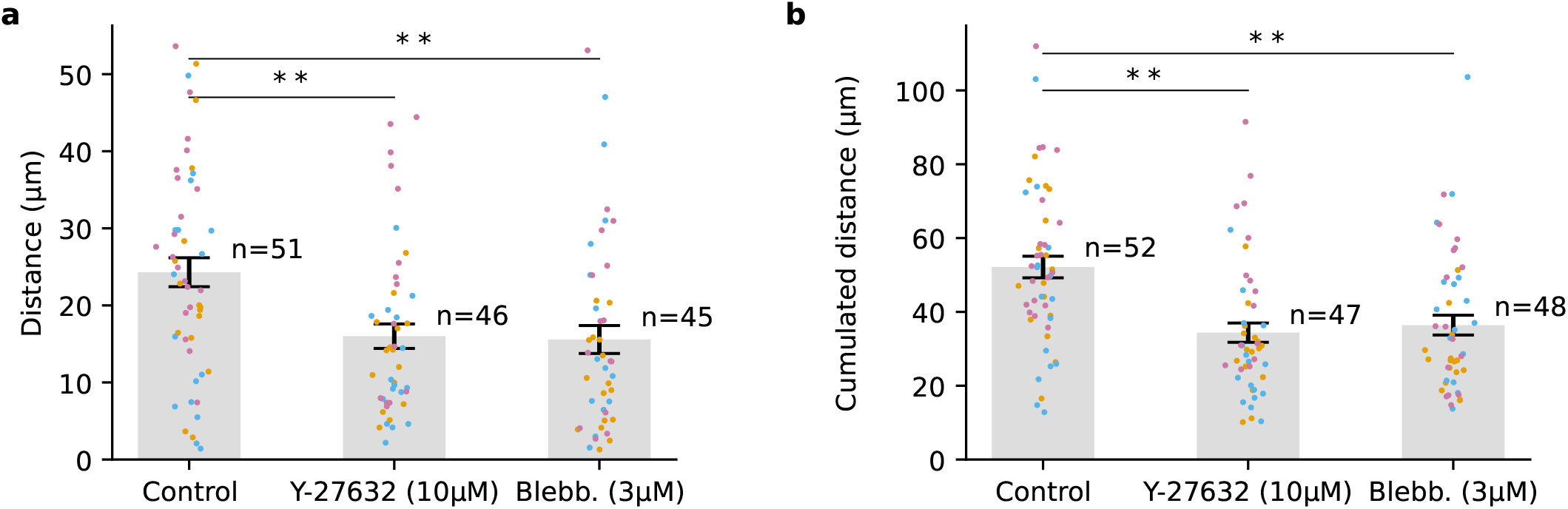
Migration distance of NK92 cells within a 23 min measurement period. a, Maximum Euclidean distance (from starting position) of cell migration after 23 min in 1.2 mg/ml collagen gels. Points show data from individual cells (from three independent experiments, indicated by color), bars show meanise; ** indicates p<0.01 for two-sided t-test **b**, Path length of cell trajectory within 23 min in 1.2 mg/ml collagen gels. Time interval between individual frames is dt= 1 min.

**Supplementary Information 6:**
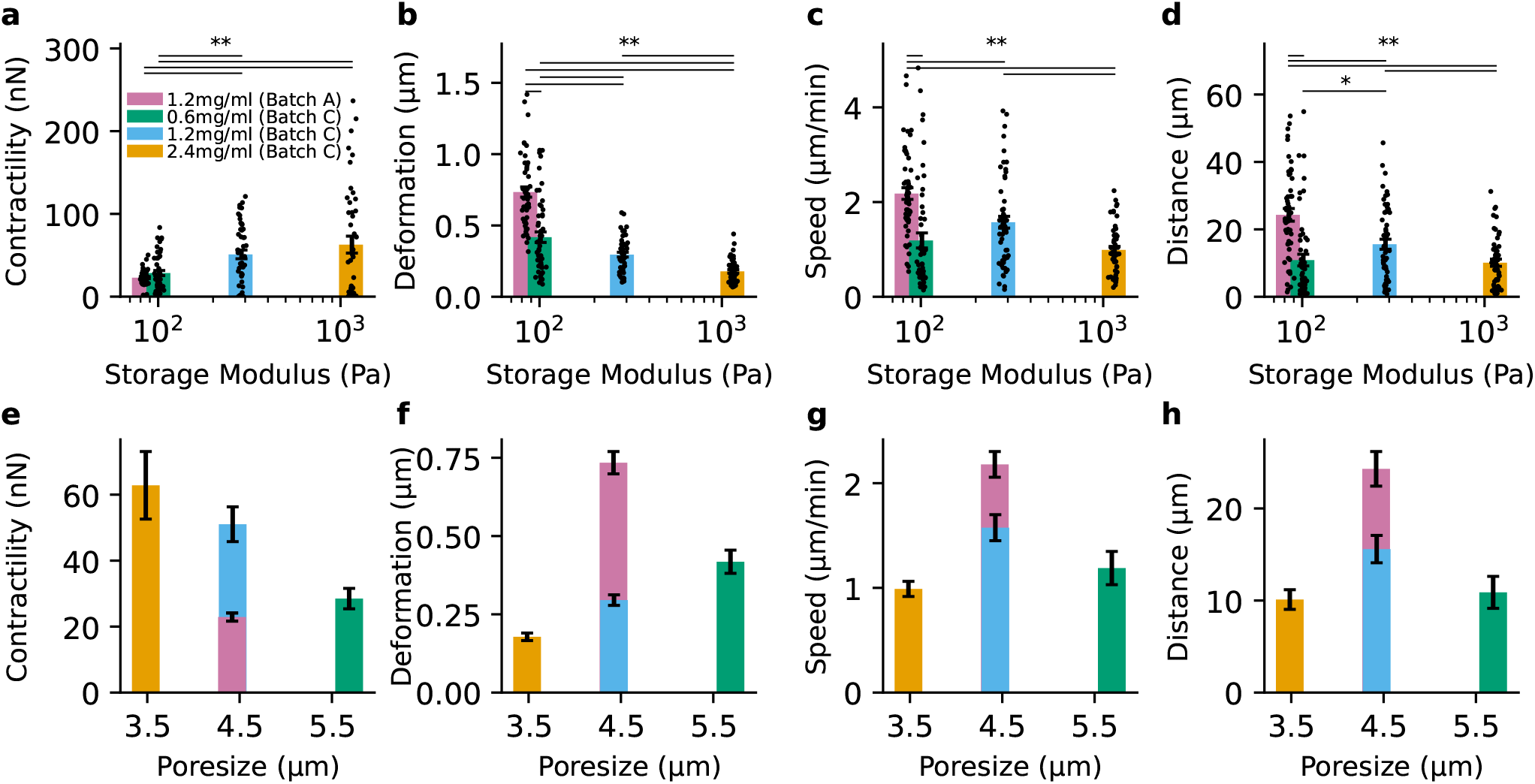
Dependence of immune cell migration and force generation on matrix stiffness and pore size. Cell contractility (**a**,**e**), matrix deformations (**b**,**f**), cell speed (**c**,**g**), and cell travelled distance (**d**,**h**) are measured in collagen gels (0.6 mg/ml, 1.2 mg/ml, and 2.4 mg/ml from different collagen batches) with different stiffnesses (**a**-**d**) and pore sizes (**e**-**h**). The storage modulus of different collagen gels is measured with a cone-plate shear-rheometer at 0.02 Hz (1% strain amplitude, see SI Fig. 3), and the pore size is derived from confocal reflection images (see SI Fig. 4). Black points indicate measurements from individual cells (total of 44-56 cells from three independent experiments), colored bars and error bars indicate mean±se, * indicates p<0.05 and ** indicates p<0.01 for two-sided t-test. (**a**,**e**), Maximum contractility of each cell during a 23 min measurement period. (**b**,**f**), Maximum of the absolute matrix deformation vector (99% percentile) of each cell during a 23 min measurement period. (**c**,**g**), Mean cell speed during a 23 min measurement period. (**d**,**h**), Maximum Euclidean migration distance from starting position of each cell during a 23 min measurement period. Cell contractility monotonically increases, and matrix deformation monotonically decreases with matrix stiffness. Migration speed and travelled distance show a maximum response at intermediate pore sizes.

**Supplementary Information 7:**
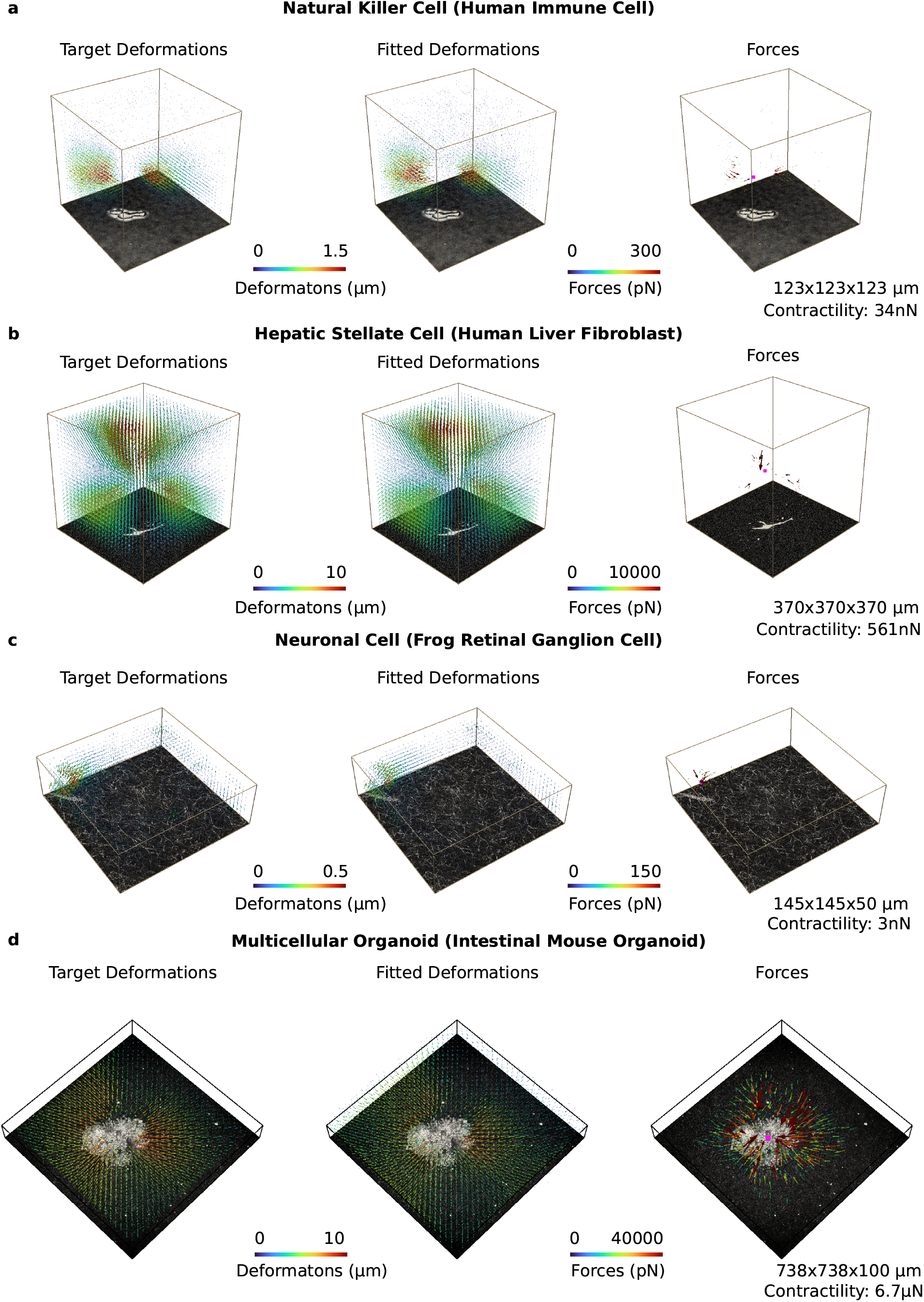
Force reconstitution for different cell types. For different cell types, cell-generated matrix deformations are measured and then interpolated onto a finite-element mesh (left). The reconstructed matrix deformations (center) and forces (right) are obtained using Saenopy. The force epicenter is shown in pink, and the size of the image stack is indicated in the lower right. **a**, Human natural killer cell (NK92) embedded in a 1.2mg/ml collagen gel (Batch A) during a contractile phase (Fig.2, SI Video 24-26). **b**, Hepatic stellate cell (human liver fibroblast) in a 1.2 mg/ml collagen gel (Batch C) after 2 days of culture (SI Video 28). **c**, Axon growth cones of a frog retinal ganglion cell embedded in a 1.0 mg/ml collagen gel (Batch D) during a contractile phase (SI Video 27). **d**, Mouse intestinal organoid in a 1.2mg/ml collagen gel (Batch C) after 24hours. Time-lapse images of organoid contraction and drug-induced relaxation are shown in SI Video 29.

**Supplementary Information 8:**
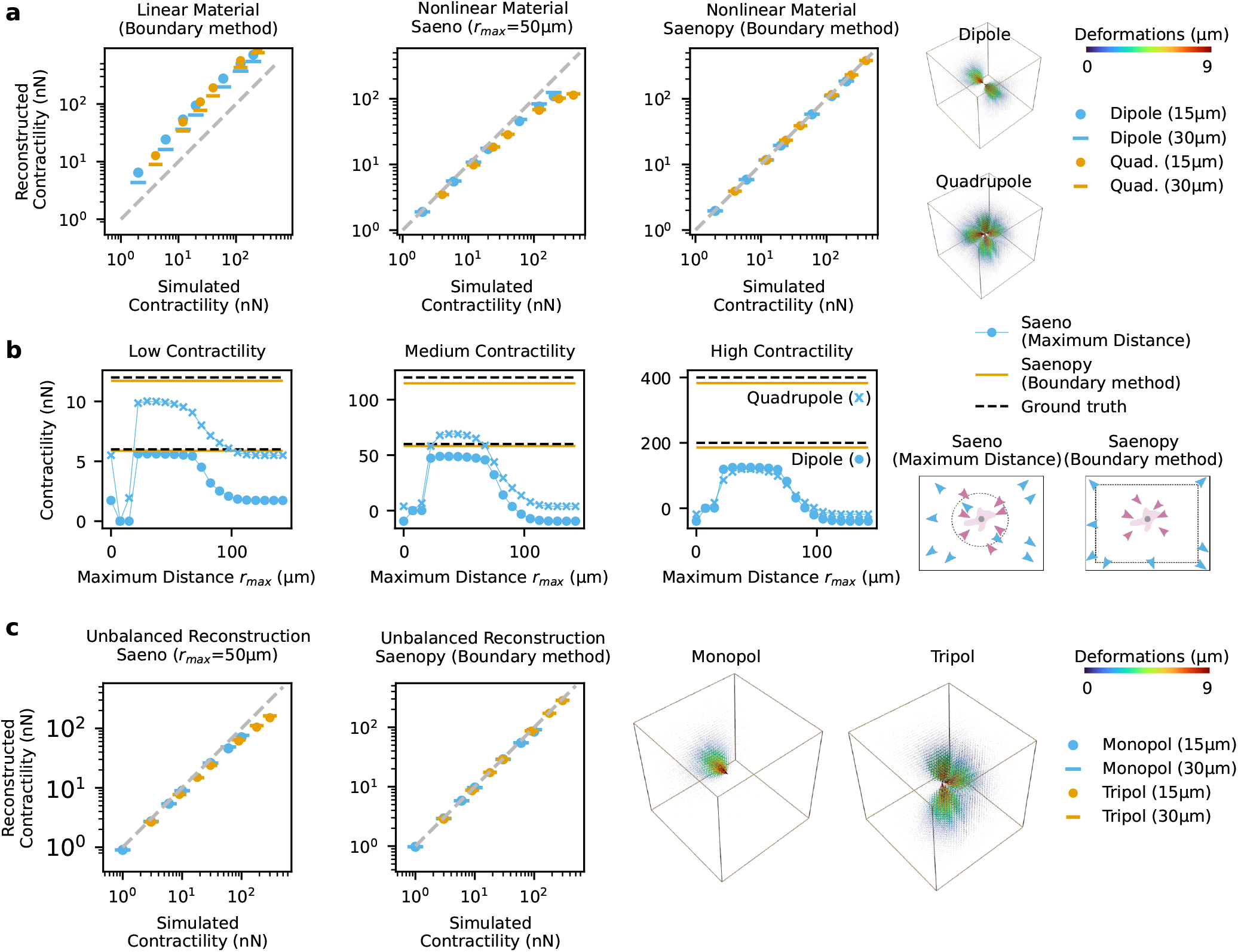
Force reconstruction benchmark tests. **a**,Benchmark tests comparing the accuracy of force reconstruction using linear Saenopy (Boundary Method, linear elastic material model without parameters *λ*_*s*_, *d*_*0*_ and *d*_*s*_, left), Saeno (Maximum Distance, *r*_max_=50 μm, middle), and non-linear Saenopy (Boundary Method, right), for artificial dipoles (blue) and quadrupoles (orange). Force reconstruction with linear Saenopy tends to overestimate total contractility. Force reconstruction with Saeno (maximum distance) tends to underestimate total contractility for high forces. Force reconstruction with non-linear Saenopy gives accurate contractility estimates over the entire simulated range. Images (right) show simulated deformation fields around a dipole (200 nN) and a quadrupole (400 nN). Reconstructed contractility for dipoles (circles) and quadrupoles (crosses) with low (6 and 12 nN), medium (60 and 120 nN) and high contractility (200 and 400 nN) versus maximum distance parameter *r*_max_ for Saeno (blue). For comparison, results from the Saenopy Boundary Method (orange) and ground-truth values (black) are shown as lines. Compared to the boundary method, Saeno shows larger deviations for all *r*_max_ values especially at high contractilities. **c**, Reconstruction of unbalanced forces from monopoles and tripoles (a single point force is removed from a dipole or quadrupole) using Saenopy (boundary method) and Saeno (maximum distance, *r*_max_=50 μm). For larger forces, Saenopy (boundary method) outperforms Saeno.

**Supplementary Information 9:**
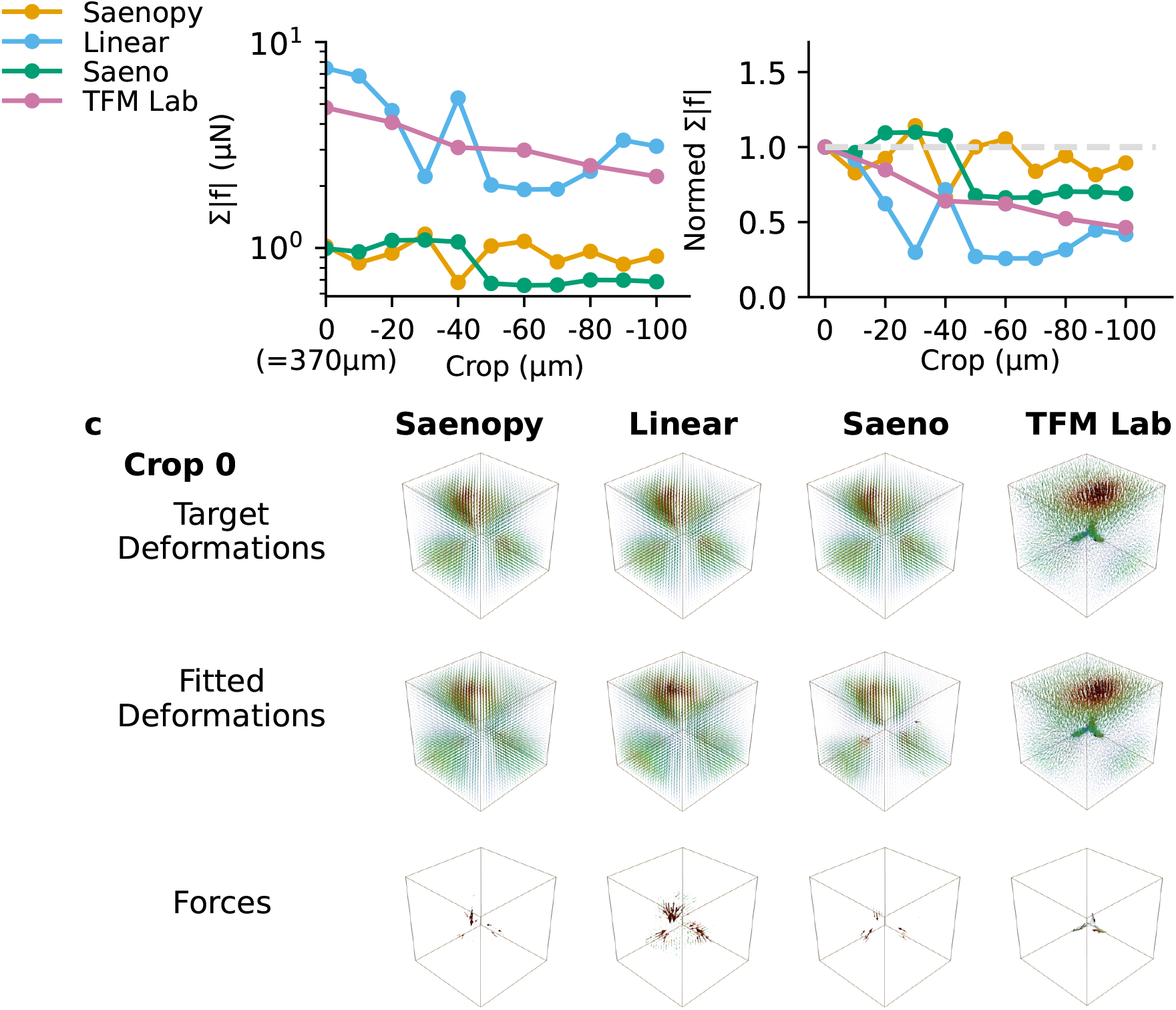
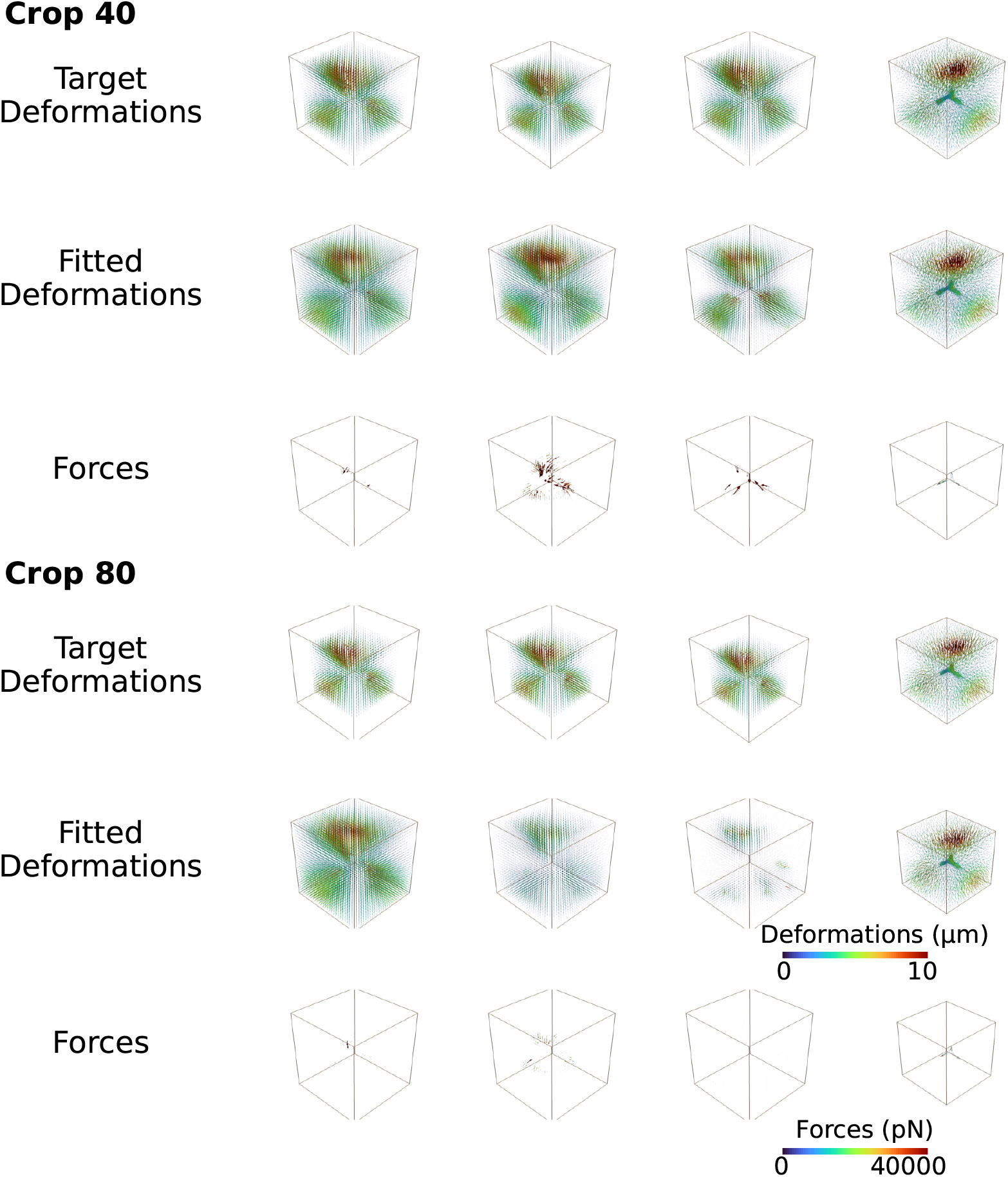
Dependence of reconstructed force on image stack size for different reconstruction methods. A representative hepatic stellate cell embedded in collagen (1.2 mg/ml, Batch C) is imaged with confocal reflection microscopy. Image stacks are cropped in x,y,z-direction by different amounts. From the cropped image stacks, we calculate matrix deformtions and reconstruct forces. We compare Saenopy (Boundary Method), linear Saenopy, Saeno (Maximum Distance method, *r*_max_ set to 75 μm), and TFM Lab software (9). For Saenopy and Saeno, we reconstruct the cell forces in a volume of (370 μm)^3^. For reconstruction with TFM Lab, we chose an elastic modulus of 1010 Pa (Batch C, 1) and a Poisson’s ratio of 0.25, and additionally provide information of the cell surface based on fluorescence images (calcein). We choose a comparable number of finite elements for all four methods. From the reconstructed force vectors, we calculate the sum of absolute forces (top left). Linear elastic material assumption (linear Saeno, and TFM Lab) results in large forces (to compensate for long-range-propagated matrix deformations that are not explained by linear material properties). Normed total forces (top right) are normalized to the respective values obtained from uncropped stacks. Reconstructed forces decrease with cropping in the case of TFM Lab and linear Saeno, and are more stable in the case of Saenopy.

**Supplementary Information 10:**
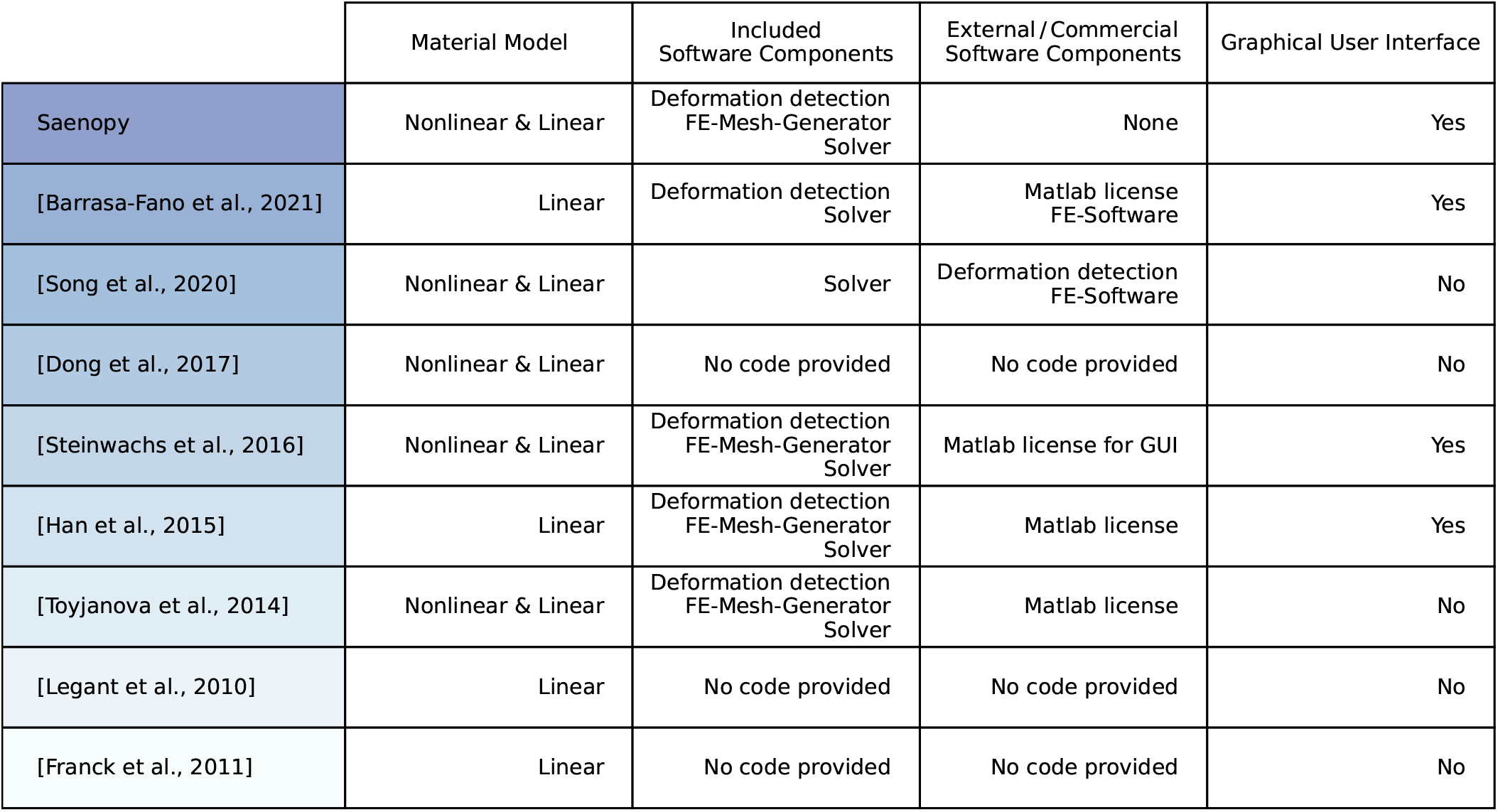
Overview of different TFM methods. Comparison of Saenopy with published 3D TFM methods regarding material models, included software components, required external or commercial software components, and graphical user interface. [Steinwachs et al., 2016 (12)], [Barrasa-Fano et al., 2021 (9)], [Song et al., 2020 (13)], [Dong et al., 2017 (14)], [Han et al., 2015 (2)], [Toyjanova et al., 2014 (15)], [Legant et al., 2010 (10)], [Franck et al., 2011 (11)].

**Supplementary Information 11:**
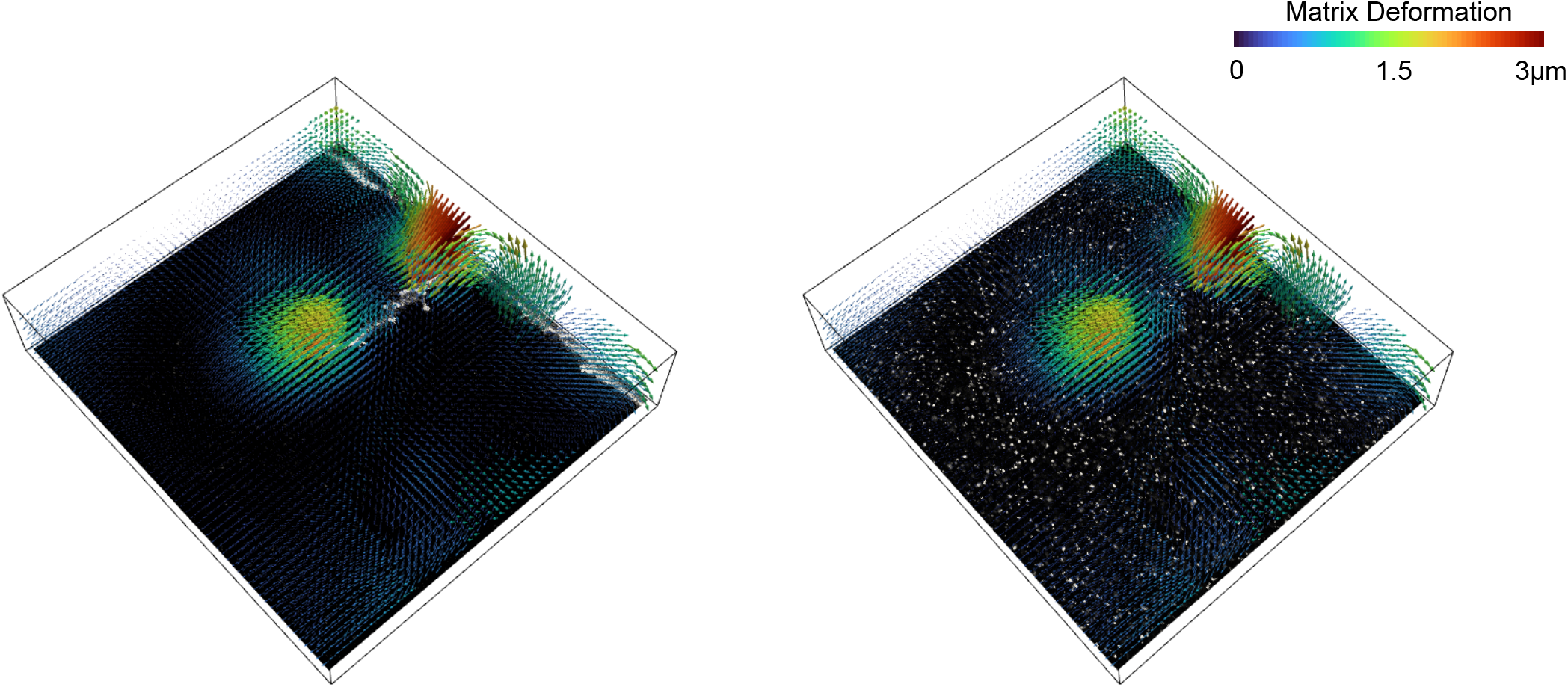
Deformation field detection using fluorescent microbeads. Matrix deformations around a human umbilical vein endothelial cell (HUVEC) in a polyethylene glycol (PEG) hydrogel (based on image stacks containing 200 nm beads, images taken from (9). Left: HUVEC cell outline; right: image of microbeads-containing PEG gel. Bounding box has dimensions of 290×262×44 μm. Matrix deformations are computed using a PIV window-size of 30 μm and an element-size of 5 μm.

**Supplementary Information 12:**
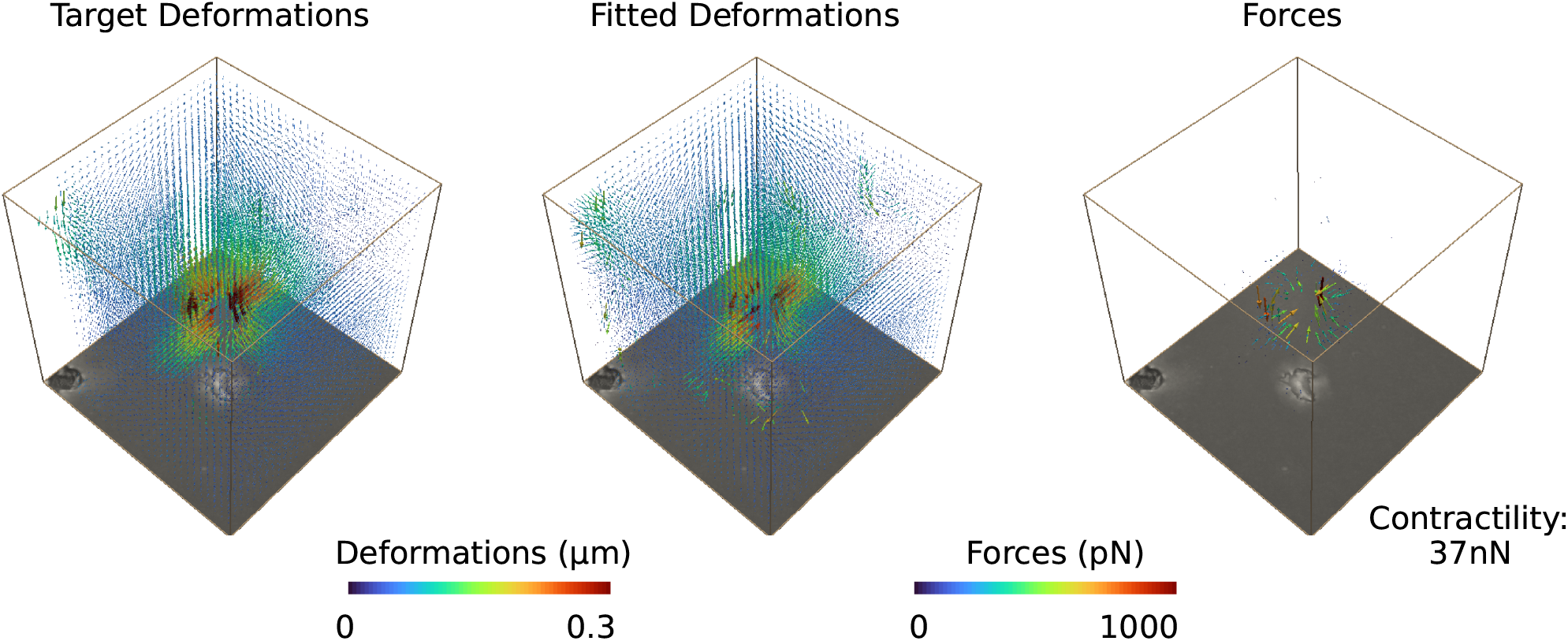
Saenopy resolves 3D force fields from bright-field image stacks. Matrix deformations and forces around a NK92 cell embedded in a 1.2 mg/ml collagen gel (Batch C). Bright-field images are acquired with an ASI RAMM microscope (Applied Scientific Instrumentation, Eugene), CMOS-camera (acA4096-30um, Basler, Ahrensburg), and 20x objective (0.7NA, air, Olympus, Tokyo). The cells are kept at 37°C and 5% CO_2_ in a stage incubator (Tokai HIT, Fujinomiya). Matrix deformations are calculated from bright-field image stacks (120 × 120 × 120 μm, with a voxel-size of 0.15 × 0.15 × 2 μm, dt = 30 s) during a contractile phase (SI Video 30, at t = 2 min). Maximum intensity projected image stacks are shown below the 3D cubes. The force reconstruction is performed using a regularization parameter of 10^11^.

**Supplementary Information 13:**
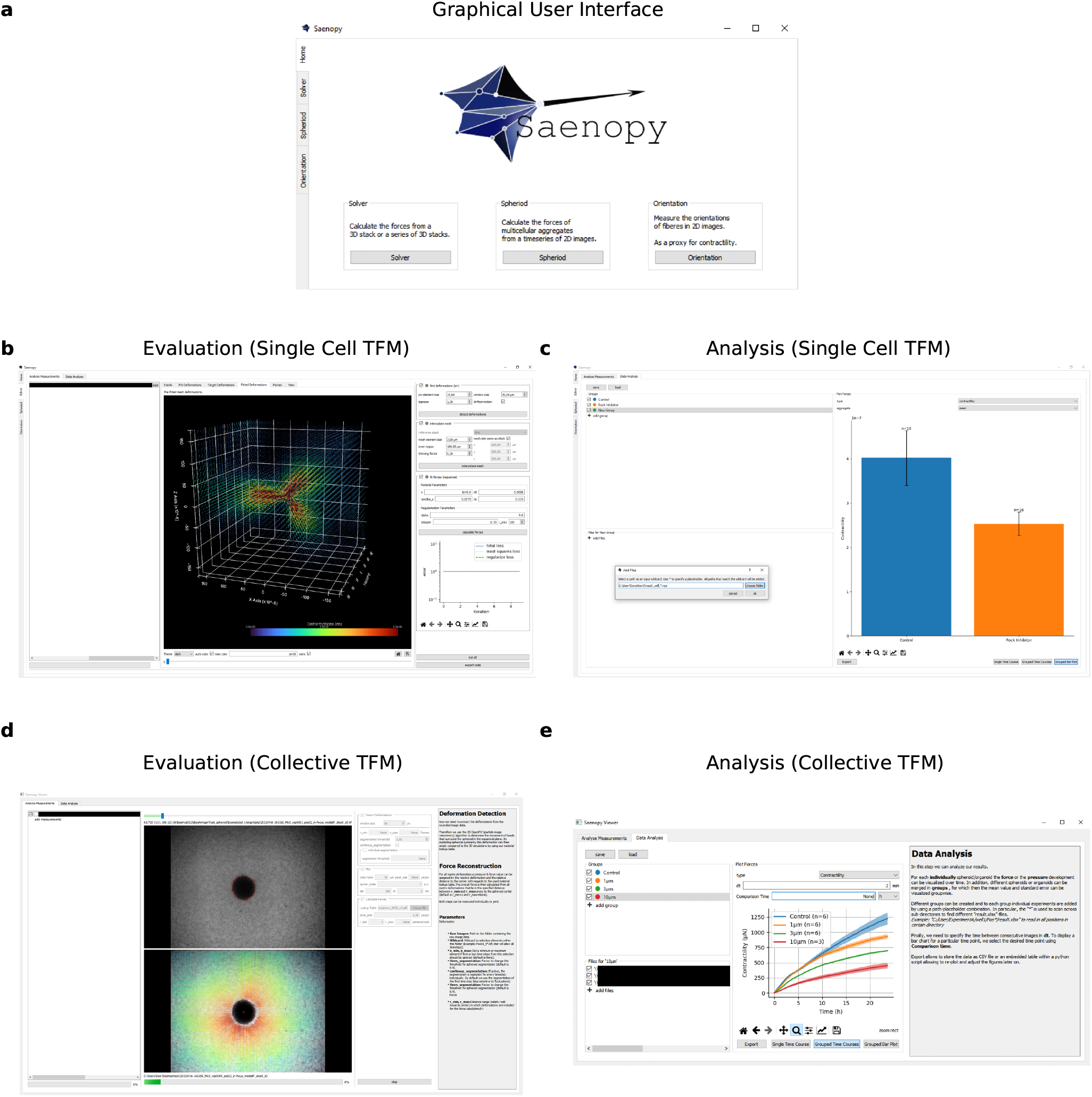
Graphical User Interface. Graphical user interface for the open-source python package Saenopy (43) for dynamic or static traction force measurements of single cells (**a**,**b**) or for multicellular aggregates (**d**: spheroids and organoids (21**?**)). Results can be evaluated and visualized individually or for grouped experiments (**c**,**e**).

**Supplementary Information 14:**
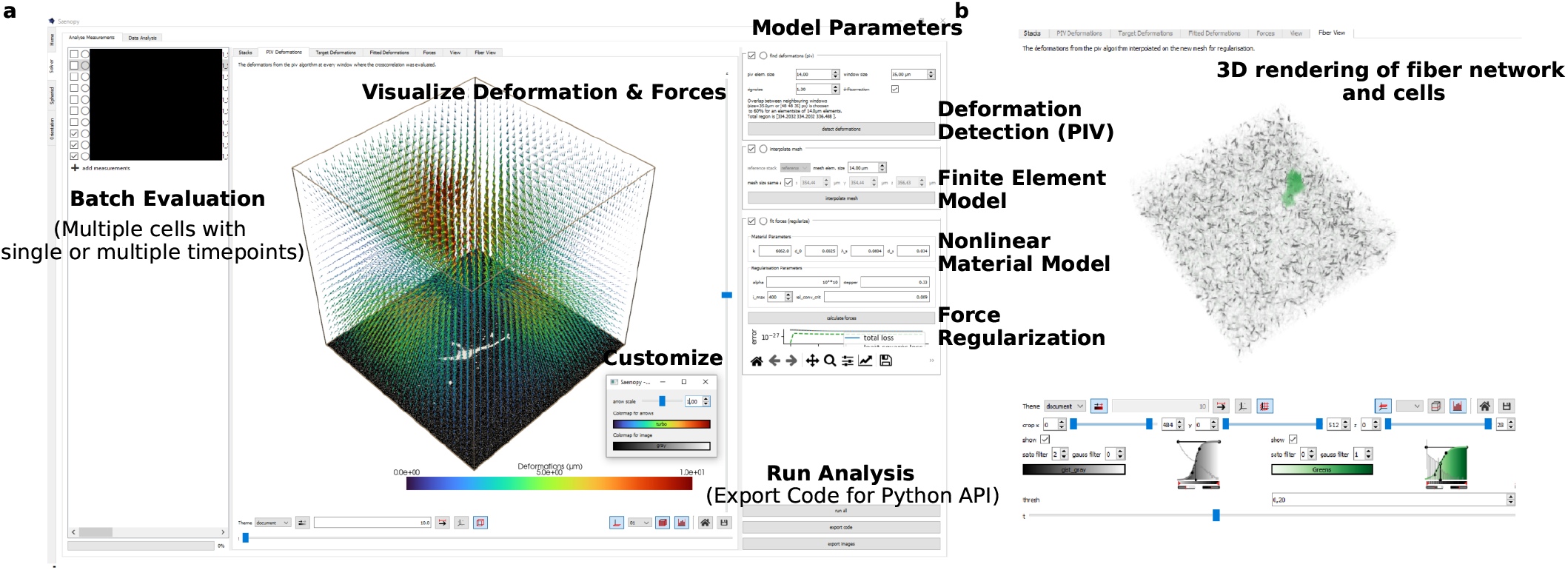
Graphical User Interface for batch evaluation. The user interface allows for flexible batch evaluation of multiple measurements, adjustment of individual analysis parameters, interactive visualisation of 3D force and deformation fields (**a**), and 3D-rendering of fiber networks and cells (**b**).

**Supplementary Information 15:**
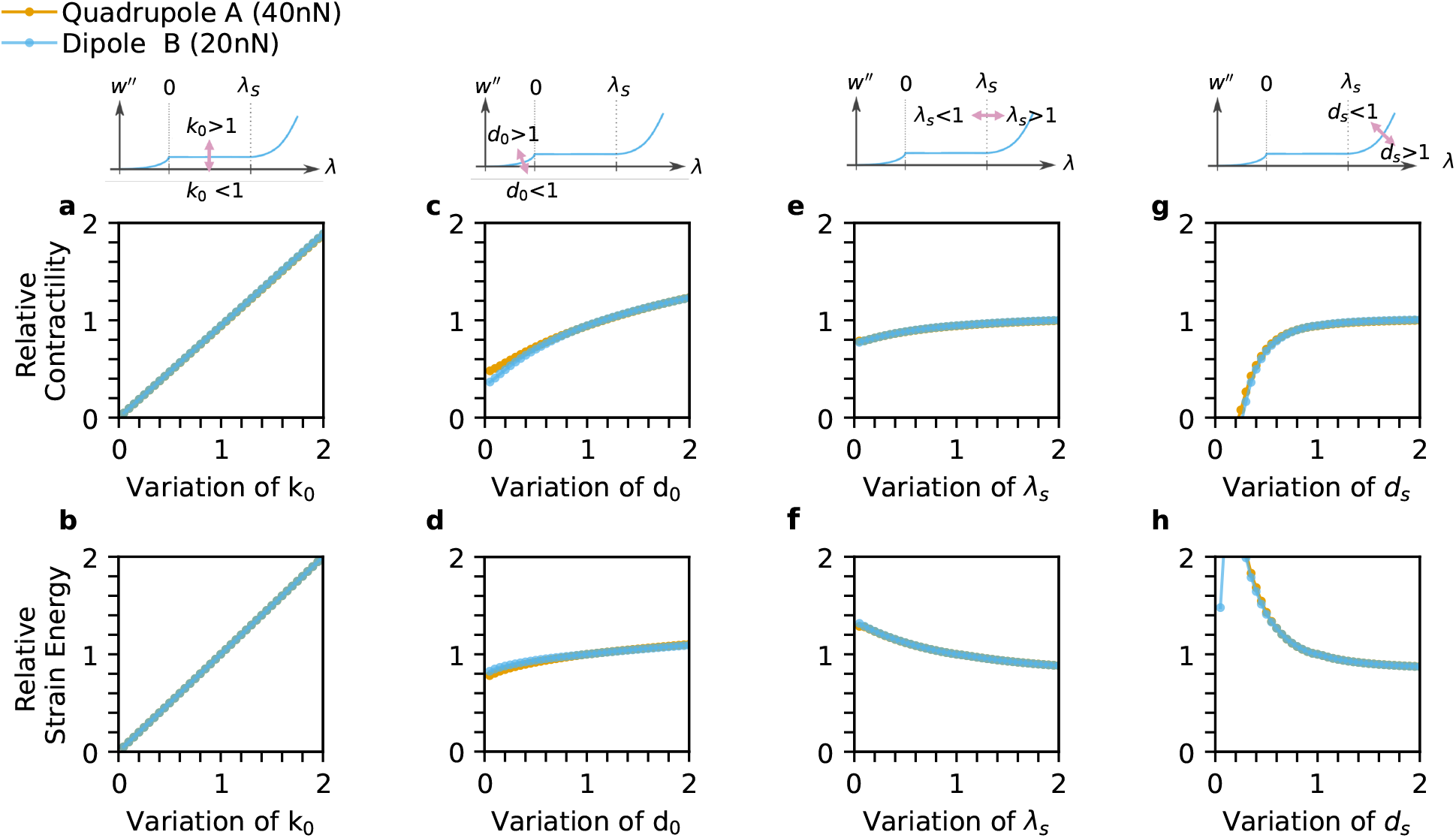
Influence of material parameters on force reconstruction for dipoles and quadrupoles. 3D matrix deformation fields are simulated for a quadrupole with a total contractile force of 40 nN (orange) and a dipole with a total contractile force 20 nN (blue) that contract in a 1.2 mg/ml collagen gel (Batch A, SI Fig. 1). We then evaluate how the reconstructed cell contractility and strain energy are affected by variations of the material parameters. As in SI Fig. 16, we alter the four material parameters (linear stiffness *k*_*0*_ (**a**,**b**), buckling coefficient *d*_*0*_ (**c**,**d**), characteristic strain *λ*_*s*_ (**e**,**f**), and stiffening coefficient *ds*) (**g**,**h**) individually with respect to the original material values used for computing the deformation field. Results are expressed as reconstructed contractility relative to ground truth of 40 nN or 20 nN, respectively, and as strain energy relative to the ground truth of the strain energy obtained for the original material values. We find that increasing linear stiffness *k*_0_ result in a proportional increase of relative contractility (**a**) and relative strain energy (**b**) regardless of total contractility. Increasing the buckling coefficient *d*_0_ results in a less than proportional increase of contractility (**c**) and strain energy (**d**). Increasing the linear strain range *λ*_*s*_ results in an weak increase of contractility (**e**) and a weak decrease in strain energy (**f**). Increasing the stiffening coefficient *d*_*s*_ results in a increase of contractility (**g**) and a decrease in strain energy (**h**). In general, we find that the contractility and strain energy for contractile quadrupoles and dipoles change proportionally, which implies that spatial arrangements of the monopole forces can be correctly inferred even with the incorrect material model.

**Supplementary Information 16:**
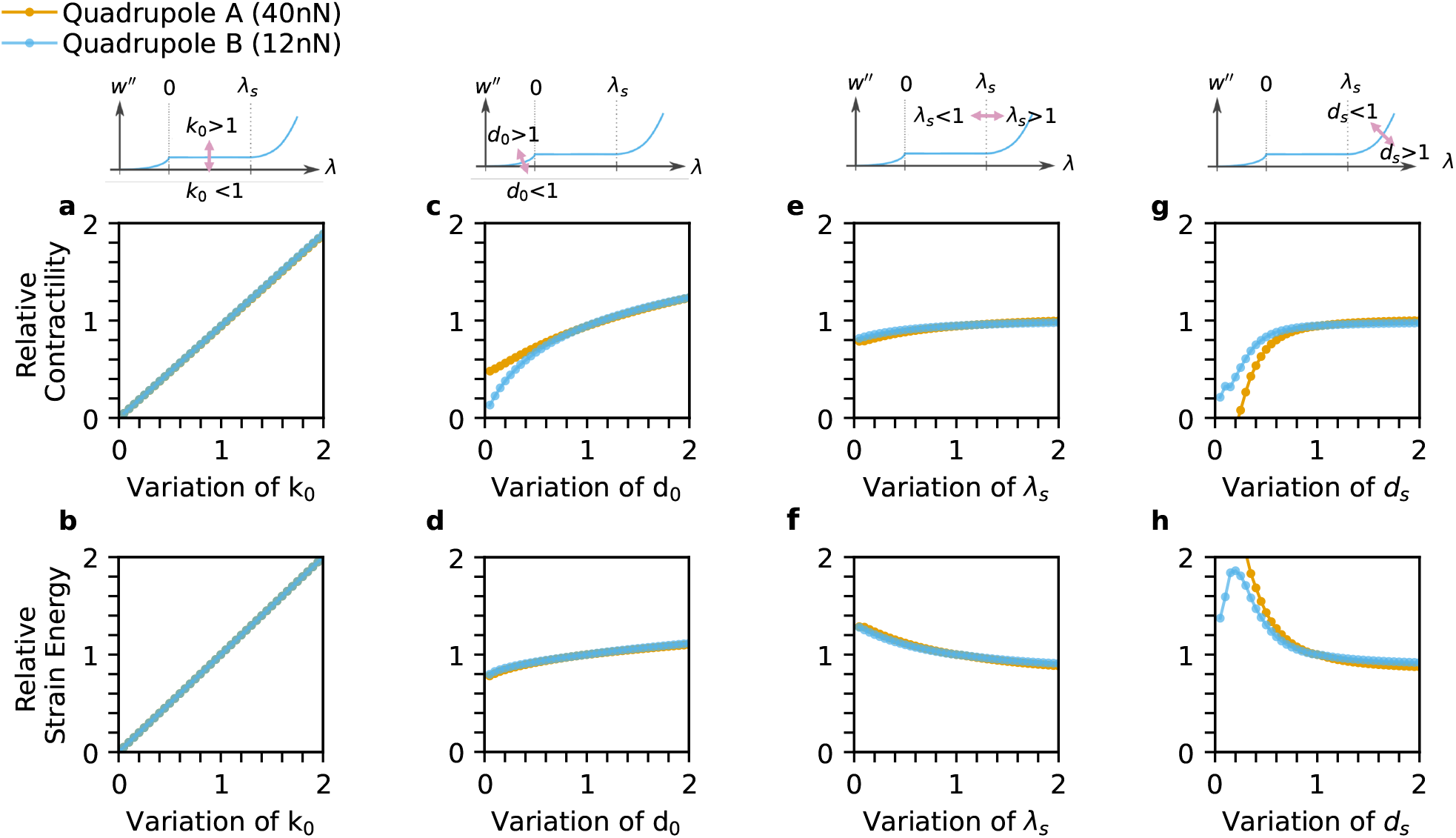
Influence of material parameters on force reconstruction for differently contractile quadrupoles. 3D matrix deformation fields are simulated for quadrupoles with a total contractile force of 40 nN (orange) or 12 nN (blue) that contract in a 1.2 mg/ml collagen gel (Batch A, SI Fig. 1). We then evaluate how the reconstructed cell contractility and strain energy are affected by variations of the material parameters. Specifically, we alter the four material parameters (linear stiffness *k*_*0*_ (**a**,**b**), buckling coefficient *d*_*0*_ (**c**,**d**), characteristic strain *λ*_*s*_ (**e**,**f**), and stiffening coefficient *d*_*s*_) (**g**,**h**) individually with respect to the original material values used for computing the deformation field. Results are expressed as reconstructed contractility relative to ground truth of 40 nN or 12 nN, respectively, and as strain energy relative to the ground truth of the strain energy obtained for the original material values. We find that increasing linear stiffness *k*_0_ results in a proportional increase of relative contractility (**a**) and relative strain energy (**b**) regardless of total quadrupole contractility. Increasing the buckling coefficient *d*_0_ results in a less than proportional increase of contractility (**c**) and strain energy (**d**), except for very low values where the contractility of the 12 nN quadrupole depends more strongly on *d*_0_. Increasing the linear strain range *λ*_*s*_ results in an weak increase of contractility (**e**) and a weak decrease in strain energy (**f**). Increasing the stiffening coefficient *d*_*s*_ results in a increase of contractility (**g**) and a decrease in strain energy (**h**). At low values of *d*_*s*_, the dependency of the 40 nN quadrupole is more pronounced. In general, we find that the contractility and strain energy for differently contractile quadrupoles change proportionally, which implies that relative changes in contractility can be correctly inferred even with the incorrect material model, except for material parameters that cause extremely non-linear material behavior (very low *d*o and *ds*).

**Supplementary Information 17:**
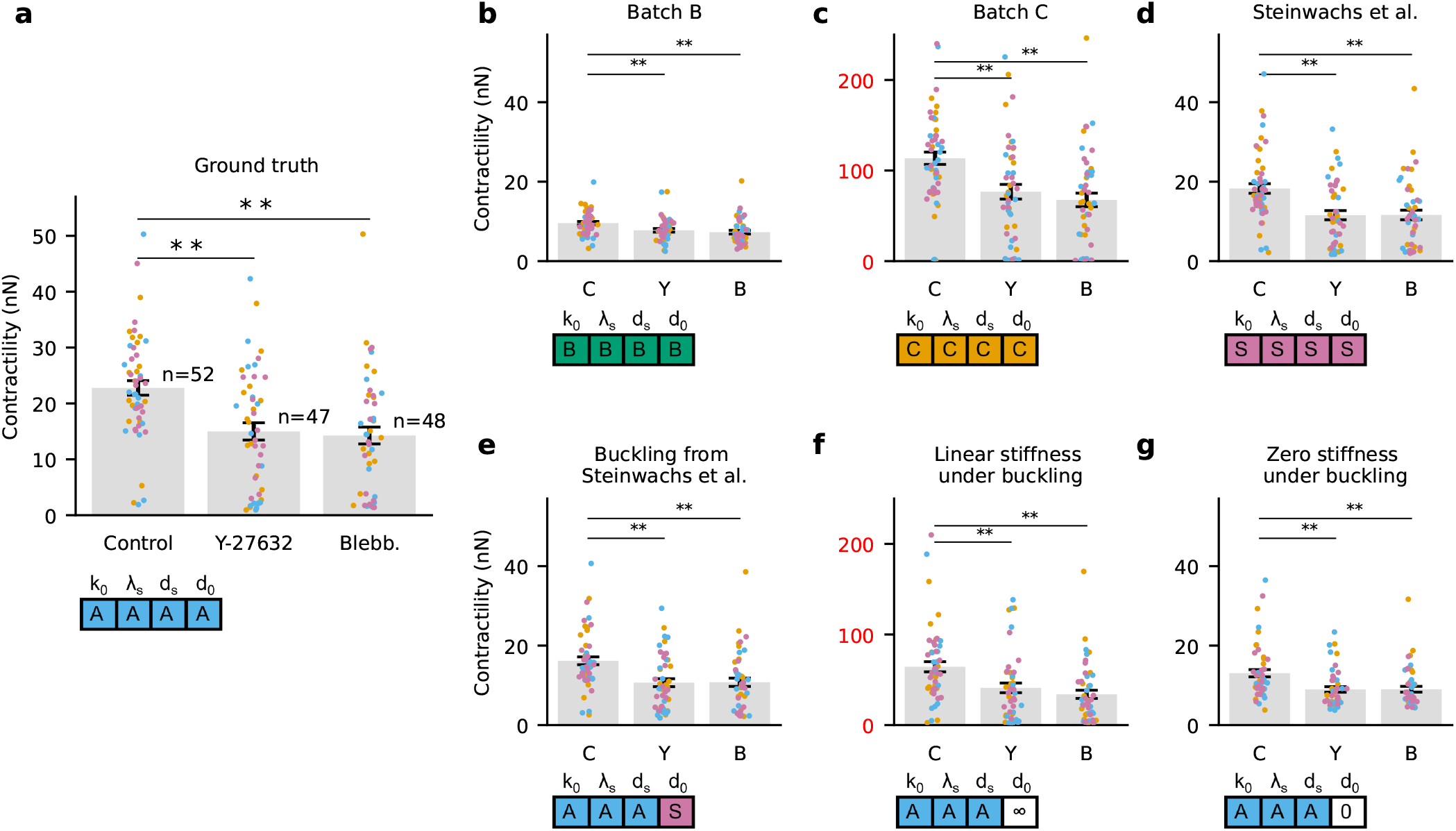
Influence of material parameters on force reconstruction for immune cells. **a**, Contractile forces (same data as shown in Fig. 2) for cells measured in 1.2mg/ml collagen of Batch A. Force reconstruction is performed with the correct material model (Batch A, “ground truth”, see SI Fig. 1). (**b-g**), Force reconstruction for the same data is performed with incorrect material models from different collagen batches: (**b**) for Batch B, (**c**) for Batch C, (**d**) for values reported in Steinwachs et al. (12), (**e**) for Batch A but with the incorrect buckling coefficient from Steinwachs et al. (12), (**f**) for Batch A but with an infinite buckling coefficient (corresponding to a linear behavior of the collagen fibers under compression), (**g**) for Batch A but with a zero buckling coefficient (corresponding to zero fiber stiffness under compression). The largest errors (highlighted in red axis labels) are observed when fiber buckling is ignored (**f**) or when the linear stiffness greatly differs from the correct value (**c**). Despite the choice of incorrect material models for force reconstruction, differences between control and drug treatment conditions remain statistically significant (p<0.01) in all cases. Points indicate data from individual cells, with independent experiments highlighted by different colors. Bars show mean±sd; ** indicates p<0.01 for two-sided t-test.

**Supplementary Information 18:**
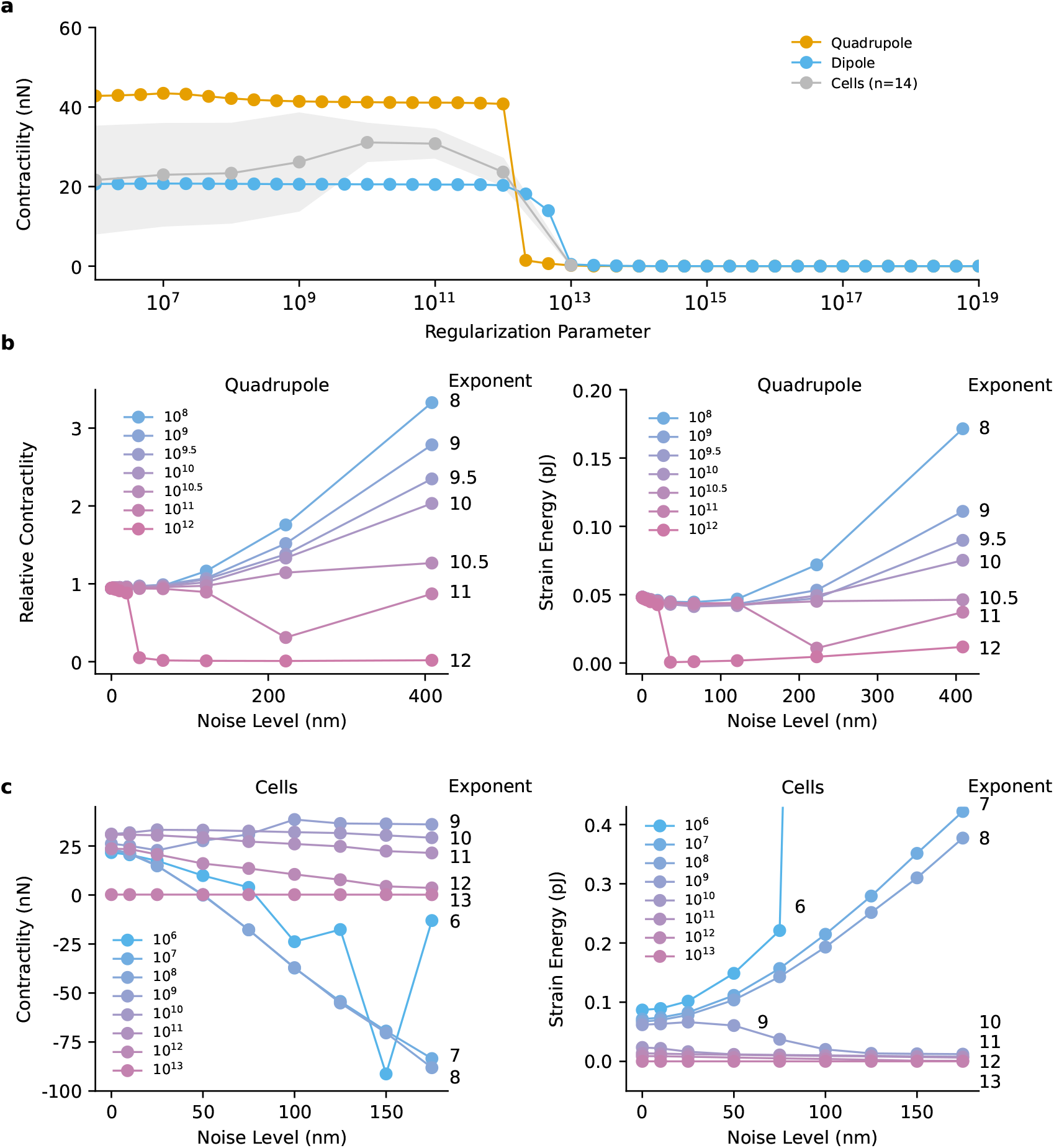
Influence of the regularization parameter on force reconstruction. **a**, Contractility of a simulated dipole (20 nN) without noise, simulated quadrupole (40 nN) without noise, and representative contractile NK92 cells (n=14, mean±se). Force reconstruction is performed with different values for the regularization parameter *α*. We find that a regularization parameter above 10^13^ leads to nearly complete force suppression. For simulated force dipoles or quadrupoles in the absence of noise, a regularization parameter of *<* 10^12^ gives the correct value. For (noisy) cell measurements, a regularization parameter around 10^10^ results in a maximum force. Lower regularization parameter can both underestimate and overestimate total cell contractility as more and more random noise forces appear, resulting in an increased variance between different cells. **b**, Contractility (left) and strain energy (right) for a simulated quadrupole (40nN) for different values of the regularization parameter and different noise levels (Gaussian noise with a specified standard deviation added to the deformation field around the quadrupole). Contractility (left) is normalized to 40 nN. In our setup, a regularization parameter between 10^10^ and 10^11^ is robust against high noise levels. Reconstructed forces are increasingly overestimated for regularization parameters below *<* 10^10^ and underestimated for regularization parameters above 10^11^. **c**, Contractility (left) and strain energy (right) for contractile NK92 cells (n = 14, standard errors are not shown for clarity). Gaussian noise on top of the experimental noise level (*σ*_*x*_*=41* nm, *σ*_*y*_ =42 nm, *σ*_*z*_ =99 nm) is added to the deformation field (as in **b**). Reconstructed forces are largely unaffected by additional noise for regularization parameters between 10^9^ and 10^11^. Smaller regularization parameters lead to increasingly erratic behavior of reconstructed forces for higher noise levels. The strain energy increases with noise for regularization parameters below 10^9^. Taken together, a choice of around 10^10^ for the regularization parameter results in the most accurate and robust force estimate in our setup.

**Supplementary Information 19:**
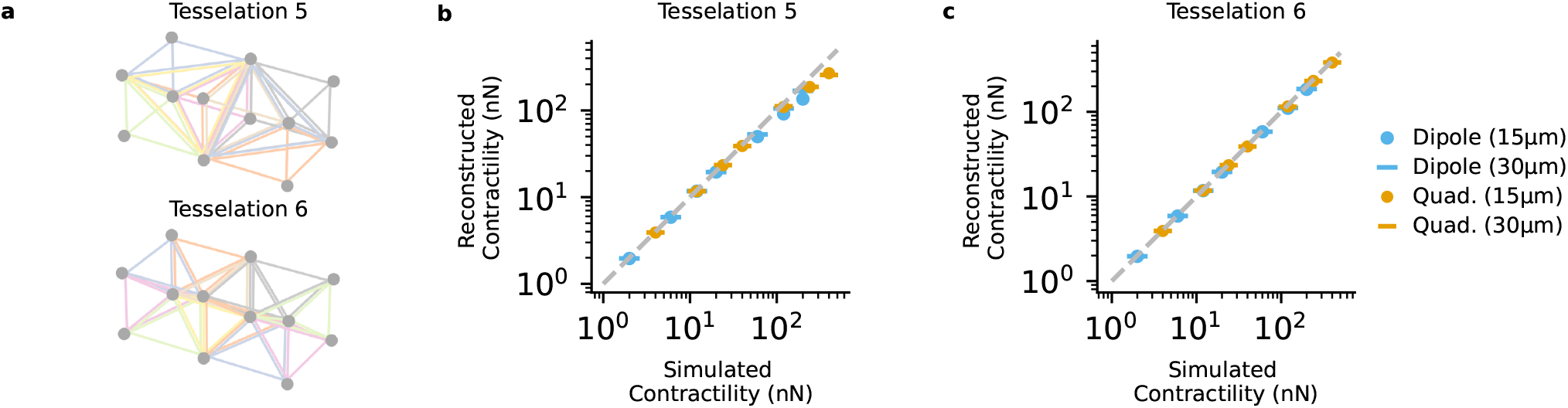
Effect of different tessellation approaches for finite-element meshes. **a**, Cubic meshes are tessellated into either 5 tetrahedra (not of equal volume) or 6 tetrahedra (of equal volume). **b**, Reconstructed contractility (for 5 tetrahedra sub-tessellation) for differently contractile dipoles (two force monopoles 15 μm or 30 μm apart) and quadrupoles (force monopoles at the vertices of a regular tetrahedron with an edge length of 15 μm or 30 μm). For total contractilities above 50 nN, the reconstructed forces become increasingly underestimated. **c**, same as in (**b**) for 6 tetrahedra sub-tessellation. In contrast to a 5 tetrahedra sub-tessellation, forces are correctly reconstructed also for higher contractilities (**c**).

**Supplementary Information 20: Video of an NK92 cell migrating through collagen (bright-field & confocal reflection)** Intensity projected image stacks of an NK92 cell migrating through a 1.2 mg/ml collagen gel (stack-size of 123×123×123 µm with a voxel-size of 0.24×0.24×1 µm) recorded every minute. The cell shows a phase of high contractility at around t = 4 min. Left: Bright-field image. Right: Confocal reflection image. Projected matrix deformations are indicated by colored arrows. Scale-bar is 20 µm. For better visualization of matrix deformations, the video is followed by a forward-backward sequence of consecutive images during a phase with high contractility.

**Supplementary Information 21: Video of an NK92 cell migrating through collagen (DIC)** Differential contrast images of an NK92 cell migrating through a 1.2 mg/ml collagen gel. Time between consecutive frames is 2 seconds. The focus has been adjusted manually during the recording. For better visualization of matrix deformations, the video is followed by a forward-backward sequence of consecutive images during a phase with high contractility.

**Supplementary Information 22: Video of an NK92 cell migrating through collagen (3D representation)**3D representation of an NK92 cell (green) migrating through dense constrictions in a 1.2 mg/ml collagen gel (brown). The cell is imaged using calcein staining (2 µM calcein AM, Thermo Fisher Scientific, Waltham) and segmented using Yen thresholding (59). Collagen fibers are imaged using confocal reflection microscopy. Images are Sato-filtered for ridge detection, highlighting the fiber structure (60). The transparency of the cell-and collagen image stacks are determined by the intensity values according to a sigmoidal transfer function. Recorded stack-size is 123x123x40 µm with a voxel-size of 0.24x0.24x1.5 µm. Time between consecutive image stacks is 1 min. Matrix deformations are indicated by colored arrows. For better visualization of matrix deformations, the video is followed by a forward-backward sequence of consecutive images during a phase with high contractility.

**Supplementary Information 23: Video of an NK92 cell migrating through collagen (3D representation)** 3D representation of an NK92 cell (green) migrating through dense constrictions in a 1.2 mg/ml collagen gel (brown). The cell is imaged using calcein staining (2 µM calcein AM, Thermo Fisher Scientific, Waltham) and segmented using Yen thresholding (59). Collagen fibers are imaged using confocal reflection microscopy. Images are Sato-filtered for ridge detection, highlighting the fiber structure (60). The transparency of the cell- and collagen image-stacks are determined by the intensity values according to a sigmoidal transfer function. Recorded stack-size is 123x123x40 µm with a voxel-size of 0.24x0.24x1.5 µm. Time between consecutive image stacks is 1 min. Matrix deformations are indicated by colored arrows.

**Supplementary Information 24: Video of matrix deformations around an NK92 cell migrating through collagen** Deformation field around an NK92 cell during migration in a 1.2 mg/ml collagen gel (stack-size of 123×123×123 µm with a voxel-size of 0.24×0.24×1 µm, dt = 1 min). The cell shows a phase of high contractility at around t = 5 min. Intensity projected bright-field image stacks are shown below the 3D fields. Note that the contractile phase occurs when the cell moves through a narrow constriction.

**Supplementary Information 25: Video of matrix deformations calculated from confocal reflection images of collagen fibers around an NK92 cell** The deformation field around the NK92 cell shown in SI Video 24 at t = 5 min (peak of the contractile phase). The deformation field is calculated from confocal reflection images using particle image velocimetry (PIV) (20). Individual images are recorded using a resonance scanner operated at 8000 Hz in combination with a galvo-stage to move the sample in z-direction. The entire image stack is recorded within 10 s (no line or frame averaging). Despite substantial image noise, the deformation field can be reliably calculated.

**Supplementary Information 26: Video showing the force reconstruction of a migrating NK92 cell** Measured matrix deformations (left), reconstructed matrix deformations (center) and reconstructed force field (right) around the NK92 cell shown in SI Video 24 at t = 5 min (peak of the contractile phase). The measured matrix deformations agree well with the reconstructed deformations. Intensity projected bright-field image stacks are shown below the 3D fields. The pink dot denotes the force-epicenter of the force field.

**Supplementary Information 27: Video of axon growth cones** Maximum projected confocal reflection image stacks of Xenopus retinal ganglion cell axon growth cones in a 1.0 mg/ml collagen gel (Batch D). Recorded stack-size is 145× 145 × 50 µm with a voxel-size of 0.14× 0.14 × 1 µm. Time between consecutive image stacks is t = 5 min. Similar to contractile phases of immune cells, we observe contractile phases during axon growth cone development. For better visualization, we show a projected volume of 124× 124 × 10 µm around the axon growth cones and show a forward-backward sequence of the contractile phases at the end of the video.

**Supplementary Information 28: Videos showing force reconstruction of a hepatic stellate cell** Confocal reflection and fluorescence image stacks around a human hepatic stellate cell in a 1.2 mg/ml collagen gels are acquired after a culture time of 2 days (stack-size of (370 µm)^3^ with a voxel-size of 0.72 ×0.72× 0.99 μm). Matrix deformation are calculated from the confocal reflection image stacks before and after relaxation of cellular forces using 10 μM cytochalsin D. The measured matrix deformation field (left) agrees well with the reconstructed deformation field (center). Intensity projected image stacks of the calcein-stained cell are shown below the 3D fields. Purple outline indicates the 3D reconstuction of the calcein-stained cell, and the pink dot represents the force epicenter (right).

**Supplementary Information 29: Video of an intestinal organoid in collagen** Image stacks of an intestinal organoid in collagen (1.2 mg/ml) are recorded using confocal reflection microscopy (stack-size=738 ×738 ×100 μm with a voxel-size of 0.74× 0.74 ×2 μm, dt = 20 min). Organoids are relaxed after 24 hours using 10 μM cytochalsin D and 0.1% Triton x-100. Images are Sato-ridge-filtered to highlight the fiber structure (60). The full time-lapse sequence is followed by a forward-backward sequence of an image pair taken before and after relaxation of cellular forces to visualize the total contraction.

**Supplementary Information 30: Video showing a contractile phase of a migrating NK92 cell measured from bright-field image stacks** Bright-field image stacks of a NK92 cell during migration in 1.2 mg/ml collagen gel (Batch C) are acquired with an ASI RAMM microscope (Applied Scientific Instrumentation, Eugene), a CMOS-camera (acA4096-30um, Basler, Ahrensburg), and a 20x objective (0.7NA, air, Olympus, Tokyo). Cells are kept at 37°C and 5% CO_2_ in a stage incubator (Tokai HIT, Fujinomiya). Matrix deformations are calculated from bright-field image stacks (dt=30 sec, volume=120× 120 ×120 μm, with a voxel-size of 0.15× 0.15 ×2 μm). Contractions are visualized by a forward-backward sequence of an image pair taken 30 s apart. Left panel show the maximum intensity projected bright-field image stack (40 μm height around the centred cell) with differential matrix deformations (arrows). The matrix deformations are clearly visible even though the individual collagen fibers cannot be resolved (right, single image plane).

