## Supplementary figures and images for "Dynamic traction force measurements of migrating immune cells in 3D matrices"

### SI_Video_24

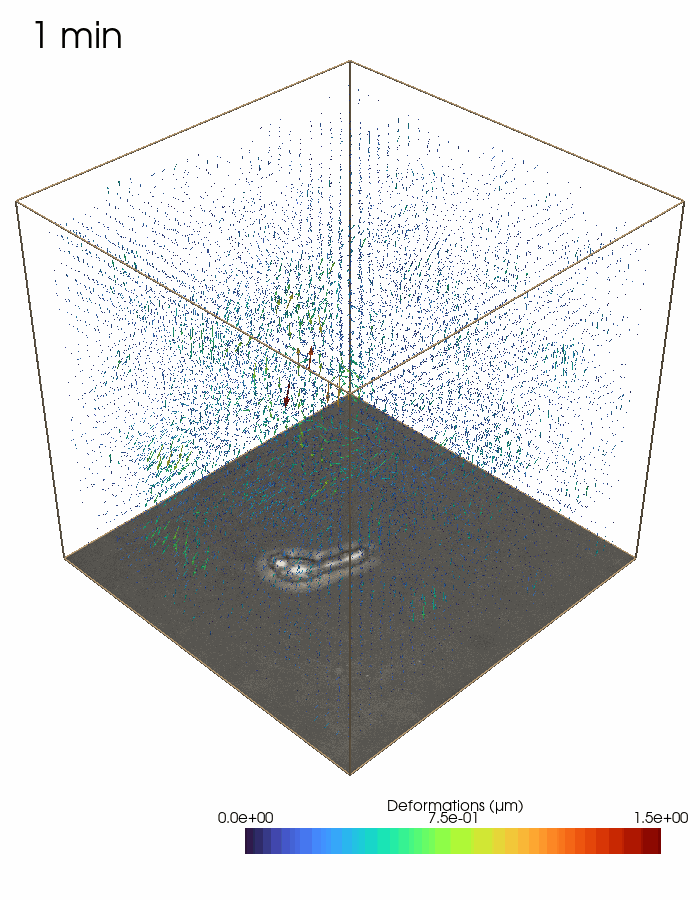

### SI_Video_25

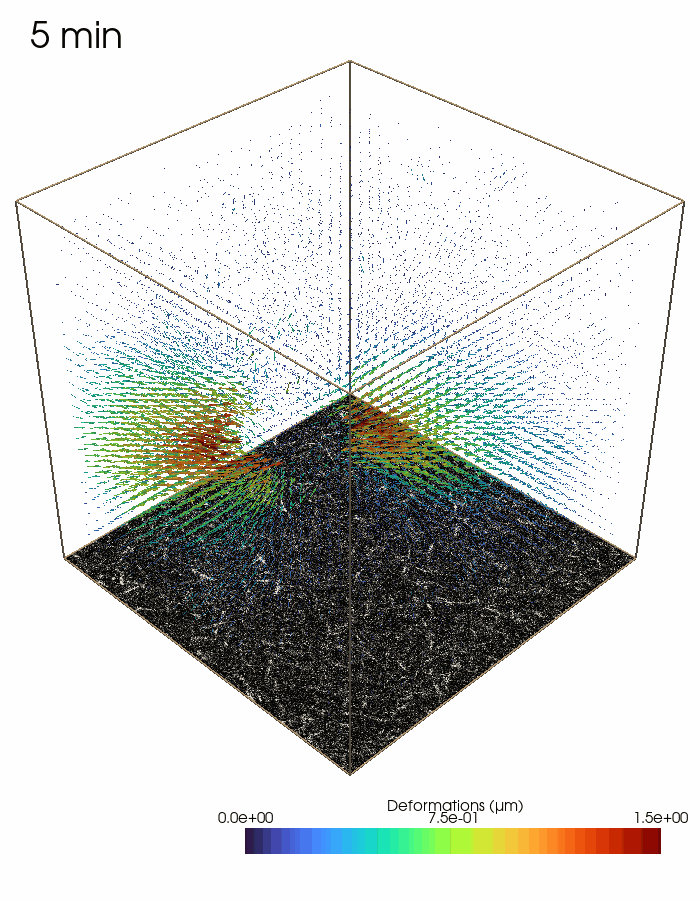
